# Spatially coordinated RTK-ERK signaling dynamics shape osteosarcoma single-cell drug response in the lung microenvironment

**DOI:** 10.64898/2026.09.30.755770

**Authors:** Rawan Makkawi, Carol Halsey, Vaibhav Murthy, Elise C. Manalo, Hugo Cros, Xin Yi, Jessie May Cartier, Matthew Chang, Mark Flory, Kelly Guptell, Amy Gross, Ryan D. Roberts, Jeremy Copperman, Alexander E. Davies

## Abstract

**Purpose:** Lung metastases are the primary cause of death in osteosarcoma patients and treatment response often vary due to convergence of genetic and phenotypic influences in the context of a dynamic tumor-host tissue environment. The purpose of this study was to determine how dynamic tumor- and host-driven receptor tyrosine kinase (RTK)-ERK signaling interactions contribute to phenotypic drug response heterogeneity.

**Experimental Design:** Live-cell time-lapse microscopy and data-driven dynamical modeling was used to resolve ERK activity, FOSL1/Fra-1 reporter expression, and spatiotemporal cell fate in living, biosensor-expressing osteosarcoma lung metastases *ex vivo* following MCL1 inhibitor exposure. RTK inhibitors were then evaluated for their ability to suppress adaptive signaling and enhance MCL1 inhibitor efficacy *ex vivo* and *in vivo*.

**Results:** MCL1 inhibition induced extensive tumor-cell death; however, persistent ERK_high_/Fra-1_high_ reporter cells emerged and were enriched at the tumor-lung interface, consistent with adaptive resistance. Experimental and computational approaches revealed that tumor-cell lysis amplified fibroblast growth factor receptor (FGFR)-mediated ERK signaling in neighboring cells, promoting a targetable drug-resistant state marked by adaptive ERK induction, FOSL1/Fra-1 expression, and correlated with upregulation of known drug resistance pathways Nrf2 and HMOX1. Combinatorial inhibition of FGFR with fexagratinib, or broad-spectrum RTK inhibitor, pazopanib, suppressed adaptive ERK activation and enhanced MCL1 inhibitor response resulting in greater efficacy *ex vivo* and *in vivo*.

**Conclusions:** Tumor-host and tumor-cell-lysis-induced FGFR-ERK signaling contribute to spatiotemporally heterogeneous adaptive resistance in metastatic osteosarcoma. These data support combinatorial targeting of RTK signaling with cytotoxic or targeted agents as a strategy to block adaptive resistance and improve treatment efficacy.

## INTRODUCTION

Lung metastases are the primary cause of death in osteosarcoma patients, with 5 year-survival rates around 20%. In patients with metastatic lesions that respond to treatment, responses to established and experimental treatments are often incomplete and/or short-duration (1,2). Patient-to-patient genetic variation is partly linked to tumor-intrinsic drug response variation; however, overall response represents the sum of genetic, phenotypic, and host tissue environment influences under conditions of treatment-induced stress over time (3–5). In the present study, we sought to understand how dynamic tissue-scale tumor-host and tumor-tumor signaling interactions contribute to adaptive drug resistance mechanisms, at single cell resolution, in metastatic osteosarcoma.

Reciprocal interactions between tumor and host cells in the metastatic lung microenvironment are increasingly recognized as major determinants of treatment response, and the establishment and outgrowth of osteosarcoma lung metastasis. Seeding of metastatic cells induces adaptive changes including wound-like responses that promote inflammatory pro-tumorigenic positive feedback loops, fibrotic stroma that often compartmentalizes or excludes immune components, and feed-forward upregulation of growth factor expression that induces protective osteosarcoma signaling programs, such as receptor tyrosine kinase (RTK)-Ras-ERK and RTK-PI3K-AKT signaling (6–9). These features are dynamically superimposed on genetic and phenotypic states, unique to each patient, thus resulting in patient-specific disease trajectories and therapeutic profiles (10–13).

Targeting of RTK-induced signaling in metastatic osteosarcoma has shown promise in clinical trials. Multi-receptor targeting drugs such as cabozantinib, regorafenib, and pazopanib have each demonstrated meaningful disease control for metastatic osteosarcoma patients (14–20). Cessation of these drugs often results in rapid disease growth, suggesting tumor control through cytostatic mechanisms, and that RTK-mediated tumor-host signaling is a mediator of tumor stability and growth but does not represent the Achilles heel of metastatic disease. Nevertheless, these data reinforce the concept that targeting tumor-host signaling interactions represents an important opportunity for suppressing metastatic lesion growth, yet improved tumor elimination will likely require a combinatorial approach, such as an RTK inhibitor-cytotoxic drug combination.

Our recent work revealed at least one mechanism by which RTK-mediated signaling promotes osteosarcoma survival and drug resistance and identified a potential combinatorial vulnerability (6). We found that host-derived signals preferentially activated tumor RTK-Ras-ERK signaling in the lung that resulted in upregulated MCL1 expression. In preclinical models bearing established metastasis, direct inhibition of MCL1 with AZD5991 produced significant antitumor activity alone and this activity was enhanced in combination with cyclophosphamide. Complementary studies have confirmed these results, reporting that MCL1 inhibition in combination with RTK inhibitors such as the broad receptor targeting inhibitor, regorafenib, or IGFR-specific inhibitor, linsitinib, were more effective than MCL1 inhibitors alone (21,22). These studies demonstrated that co-targeting of host-derived growth factors, and the tumor intrinsic vulnerabilities induced by them, could hold promise for improved osteosarcoma treatment. However, these combinations were not curative, indicating some level of heterogeneity capable of promoting fixed or adaptive resistance.

Our previous study found evidence that although host-derived factors induced RTK-ERK signaling and downstream ERK target expression, signaling and target expression was heterogenous within and between lesions. For example, ERK target genes such as MCL1 and AP-1 transcription factor, FOSL1/Fra-1, were heterogeneously expressed in both preclinical and human lung lesions, suggesting spatiotemporal variation in signaling that could impact single cell drug response and resistance (6). ERK signaling and associated downstream effectors are well known to modulate drug response through a wide variety of mechanisms (23–26). Furthermore, signaling though RTKs can produce highly dynamic ERK signaling patterns, leading to rapid temporal and spatial fluctuations in signaling amplitude and frequency that vary from cell-to-cell, and despite the presence of uniform ligand abundance in the system (27–29). The result of this variation is RTK-ERK driven cell-to-cell gene expression heterogeneity and phenotypic plasticity (27,28). Thus, in tissues with non-uniform cellular composition and ligand distribution, such as the metastatic osteosarcoma microenvironment, the potential for gene expression and phenotypic heterogeneity is amplified, which could represent a mechanism by which spatiotemporal drug response variation and/or adaptive resistance can arise.

Here, to test the hypothesis that dynamic tissue-scale signaling interactions drive tumor heterogeneity and adaptive resistance, we used Lung Serial Imaging of Tumor and microEnvironment (LungSITE), a precision cut lung slice (PCLS)-based platform coupled to biosensors and computation, (30) to track ERK signaling, FOSL1/Fra-1 expression, and cell fate within living osteosarcoma lung metastases *ex vivo*. ERK, Fra-1 (FIRE) biosensor expressing osteosarcoma cell lines were used to generate experimental metastasis and track the relationship between single-cell spatiotemporally resolved cell signaling and cell fate under MCL1 inhibitor challenge. We found that MCL1 inhibition induced extensive cell death but left a population of persistent ERK_high_/FIRE_high_ cells enriched near the tumor-lung interface. Mechanistically, we found that tumor-cell lysis amplified FGFR-mediated ERK signaling cell in neighboring cells, promoting acquisition of a drug-tolerant state correlated with high Fra-1, and expression of known chemotherapy resistance factors such as NRF2-driven stress response protein HMOX1, which were also found to be highly upregulated near the tumor-lung interface in heavily pretreated patient samples. Selective and broad RTK inhibition, including fexagratinib and pazopanib, blocked adaptive ERK signaling activation and enhanced MCL1 inhibitor efficacy *ex vivo*; and improved disease control in preclinical models *in vivo*. Together, these findings identify adaptive microenvironmental and death-induced FGFR–ERK signaling as drivers of spatiotemporally heterogeneous resistance and support the role of combinatorial RTK inhibition for metastatic osteosarcoma treatment.

## MATERIALS AND METHODS

### Cell culture and reporter cell lines

OS-17 cells were derived from an early-passage culture established from an OS-17 patient-derived xenograft obtained from a primary femur biopsy at St. Jude Children’s Research Hospital and maintained in RPMI 1640 (Gibco, 11875093) supplemented with 10% FBS (31). OS052 and OS186 cells, obtained from the Sweet-Cordero laboratory (UCSF), were maintained in DMEM (Gibco, 11995-065) supplemented with 10% bovine growth serum (Cytiva, SH30541.03)(32). The 143B cell line was obtained from ATCC and maintained in DMEM supplemented with 10% FBS.

OS-17 cells were transduced with lentiviral vectors encoding the ERK kinase translocation reporter fused to Clover, the Fra-1–based Integrative Reporter of ERK (FIRE) fused to mTurquoise, and histone H2B fused to iRFP670, each cloned into the pLJM1 backbone (27,33,34). Reporter-positive cells were selected with puromycin (1 μg/mL; Gibco, A11138-03) and isolated by flow cytometry. Reporter derivatives of 143B, OS052, and OS186 cells were generated by lentiviral transduction of ERKTR-iRFP713 and H2B-mCherry2, followed by puromycin selection. OS-17-FLUC cells were generated by lentiviral transduction of firefly luciferase as previously described (6). OS-17-H2B lines expressing H2B fused to mVenus, iRFP670, mTurquoise, or mCherry were generated by lentiviral transduction and selected with geneticin (200 μg/mL; Gibco, 10131035).

### Experimental metastasis and precision-cut lung slices

All animal studies used 8- to 10-week-old NOD.Cg-*Prkdc^scid^*/J mice and were performed in accordance with institutional guidelines. For experimental metastasis, 1 × 10^6^ OS-17 or 143B reporter cells were injected into the tail vein of anesthetized mice. For clonality experiments, 1.5 × 10^5^ cells from each OS-17-H2B fluorescent reporter line were mixed to a total of 1 × 10^6^ cells before injection. Mice were euthanized 2 to 3 weeks after injection unless otherwise indicated.

For precision-cut lung slices (PCLS), lungs were inflated in situ with 2.5% low-melting-point agarose prepared in phenol-free DMEM/F12, excised, and maintained on ice. Lungs were then cultured in DMEM/F-12 supplemented with insulin (250 ng/ml, Sigma-Aldrich, I1882), transferrin (10 µg/ml, Sigma-Aldrich, T8158), β-estradiol (10^-10^ M, Sigma-Aldrich, E2758), hydrocortisol (1.4 × 10^-6^ M, Sigma-Aldrich, H4001), sodium selenite (2.6 ng/ml, Sigma-Aldrich, S5261), vitamin C (0.1 µg/mL, Sigma-Aldrich, A4403), vitamin A (0.2 mg/mL, Sigma-Aldrich, R2625) and primocin (100 µg/ml, InvivoGen, Ant-pm-1). Media was changed every other day.

### Live-cell and live-tissue imaging

Imaging was performed using a Nikon Ti2 automated microscope equipped with an environmental chamber maintained at 37°C and 5% CO2, SOLA II LED illumination, an Orca Fusion BT camera, and NIS-Elements AR software. For monoculture experiments, collagen-coated glass-bottom 96-well plates were seeded with 7,000 cells per well and incubated overnight. Cells were switched to phenol-free DMEM/F12 imaging medium containing transferrin, hydrocortisone, β-estradiol, and Hoechst 33342 and equilibrated for 4 hours before imaging. Images were acquired every 6 minutes.

For PCLS imaging, slices were transferred to glass-bottom 24-well plates and secured with custom acrylic clamps. Slices were washed twice in phenol-free DMEM/F12 and cultured in imaging medium containing transferrin, hydrocortisone, β-estradiol, vitamins C and A, primocin, and Hoechst 33342. Unless otherwise specified, treatments were added approximately 6 hours before imaging; AZD5991 was added at the beginning of imaging. Images were acquired every 20 or 30 minutes for 20 to 48 hours. For longer experiments, medium and treatments were replenished every other day. Independent replicate experiments used PCLS obtained from different mice.

Treatments included FGF2 (100 ng/mL), PD0325901 (100 nM), MK-2206 (1 μM), AZD5991 (100-500nM), or fexagratinib, NVP-AEW541, erlotinib, CP-673451, pazopanib, dovitinib, erdafitinib, or nintedanib at 2μM, and cell-line-derived lysates (10 μg per well). AZD5991 concentrations used in subsequent experiments were selected from independent dose-response assays performed for each inhibitor vial. The lower concentration corresponded to the OS-17 cell-viability IC50 (100 nM), and the high-dose condition was five-fold higher (500 nM).

### In vivo drug efficacy studies

For therapeutic studies, mice were inoculated intravenously with 1 × 10^6^ OS-17-FLUC cells and monitored weekly by bioluminescence imaging. Mice with established pulmonary metastases were randomized to vehicle, AZD5991, pazopanib, or combination treatment. AZD5991 was administered intraperitoneally at 60 mg/kg three times weekly. Pazopanib was administered orally at 30 mg/kg twice daily. Treatment continued for 2 weeks or until humane endpoints were reached. Body weight and clinical condition were monitored twice weekly. Lungs were perfused with saline and collected for formalin fixation.

### Image analysis

2-dimensional live-cell and immunohistochemical images were analyzed using MATLAB R2025a and custom processing code. Nuclear fluorescence or Hoechst staining was used for segmentation and tracking. ERK-KTR cytoplasmic-to-nuclear fluorescence ratios, FIRE expression, and antibody fluorescence intensities were quantified over time. At least two independent replicate experiments and approximately 6,000 cells per condition were analyzed where applicable.

PCLS datasets (**Supplementary Tables 1 and 2**) were analyzed using the Serial Imaging of Tumor and microEnvironment (SITE) pipeline (30) based in Python. Tumor-cell cytoplasmic and nuclear masks were generated using fine-tuned Cellpose models trained on manually annotated images. Cells were linked over time using the SITE optimal-transport tracking framework. Segmentation and tracking were manually validated on representative images.

ERK activity was quantified as the cytoplasmic-to-nuclear ERK-KTR intensity ratio, whereas FIRE activity was measured as mean nuclear fluorescence. Death events were identified from characteristic morphological changes and manually validated over a 2-hour trajectory endpoint window. Death-to-ERK signaling correlations were calculated as a function of distance and time from an annotated cell death event, with shuffled death-event assignments used as null controls.

Local tumor–microenvironment organization was quantified from 3D cell, lung-tissue, and extracellular boundary maps. Inverse-distance proximity features were calculated at 1-μm boundary resolution and summarized using spherical multipole expansions. The isotropic (l = 0) and first anisotropic (l = 1) components of cancer-cell and host-tissue proximity were used to define three contact-niche states: low cancer proximity (LCP), mixed cancer proximity (MCP), and high cancer proximity (HCP).

Recent exposure to cell death was quantified by summing inverse-distance-weighted contributions from death events occurring within the preceding 8 hours. ERK-KTR states were binarized at a cytoplasmic-to-nuclear ratio of 1.0, and OFF→ON and ON→OFF transition probabilities were calculated as a function of death proximity.

For adaptive drug-response modeling, each trajectory step was assigned to one of 24 composite states defined by contact niche, viability, death-proximity state, and ERK state. Empirical transition matrices were estimated from consecutive trajectory steps. Death rates were estimated separately for ERK-KTR_high_ and ERK-KTR_low_ cells and incorporated into the transition matrix; dead states were treated as absorbing. Model predictions were compared with observed state occupancies over time. An in silico ERK-inhibition model was generated by blocking transitions from ERK-KTR_low_ to ERK-KTR_high_ states and renormalizing transition probabilities. Uncertainty was estimated by bootstrap resampling of imaging sets or fields of view, as specified for each analysis.

***See extended methods for a full description of experimental and computational methods***

## RESULTS

### LungSITE modeling for dynamic live tracking of metastatic osteosarcoma cell signaling and fate heterogeneity *ex vivo*

Using the Lung Serial Imaging of Tumor and microEnvironment (LungSITE) platform (**Figure 1A**), a high resolution variant of the PuMA model, we sought a more nuanced understanding of how live single cell ERK signaling and ERK effector expression heterogeneity correlate with single cell spatial and temporal drug response within the osteosarcoma lung microenvironment (30,35). To this end, we generated a set of OS-17 reporter cell lines and validated them in vitro. Lines were engineered to co-express fluorescently tagged ERK kinase translocation reporter (ERK-KTR), a degron-based reporter Fra-1-based Integrative Reporter of ERK (hereafter ‘FIRE’) for downstream ERK target gene and AP-1 transcription factor, FOSL1/Fra-1, and histone H2B to enable cell segmentation and tracking over time (**Figure 1A**)(29,33,36). We then validated reporter lines cultured directly on imaging plates by adding FGF2 or MEK inhibitor (MEKi; PD0325901) to stimulate or block ERK signaling, respectively. (**Figure 1B**, left panel). As expected, reporter cells exhibited no significant ERK reporter activity at baseline whereas FGF2 addition led to potent activation of ERK-KTR and a corresponding monotonic increase in FIRE over time (**Figure 1B)**. Interestingly, we observed a temporal delay (∼7 hours) in FIRE accumulation following ERK stimulation (**Figure 1B**, right panel), indicating that OS-17 integrate ERK activity to FIRE stabilization more slowly than described for other cell types (29). Addition of MEKi produced a modest decrease in ERK-KTR activity and FIRE signal over time, consistent with previously reported baseline low to undetectable ERK activity in OS-17 cell lines in absence of exogenous stimuli (**Figure 1B**) (6). These data confirmed performance of OS-17 reporter cells to appropriately sense and respond to exogeneous stimuli and perturbations.

**Figure 1.**
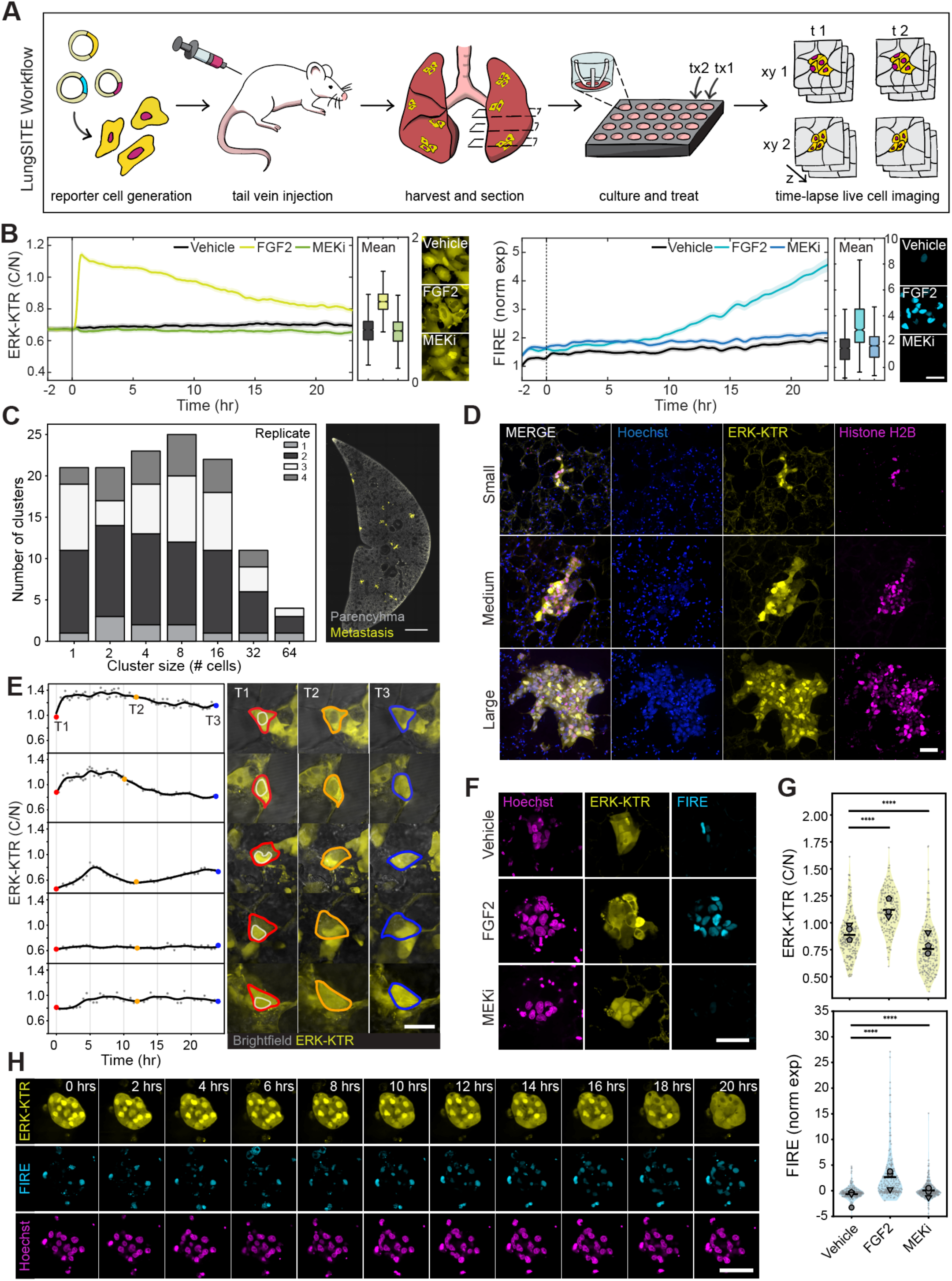
LungSITE workflow overview and validation for studying osteosarcoma cell signaling and fate *ex vivo*. (A) Schematic of LungSITE workflow depicting reporter cell generation, inoculation, tissue processing and imaging (left to right). (B) Mean traces (bold line) and inter-quartile range (shading surrounding mean) of ERK-KTR activity and normalized FIRE expression obtained from OS-17 cells via live cell imaging for the indicated durations. Box plots and images depict snapshot ERK-KTR activity immediately after treatment (t=0) and mean FIRE expression at experiment endpoint (t=24hrs). Two independent replicate experiments were performed with ≥3000 cells per condition for each experiment. Scale bar = 50 µm. (C) Stackplot (left) illustrates the size distribution range of metastases at 3-weeks post injection over four independent replicate experiments with a representative image (right) of OS-17 reporter cell lung metastases (yellow). Scale bar = 1 mm. (D) Representative 40x magnification images of small (top), medium (middle) and large (bottom) metastasis co-expressing ERK-KTR (yellow), and H2B reporter (magenta). Lung nuclei are counterstained with Hoechst (blue). Scale bar = 50 µm. (E) Representative single cell ERK-KTR activity traces, reported as cytoplasmic to nuclear (C/N) fluorescence ratios, obtained from live cell imaging of OS-17 metastasis in LungSITE for the indicated duration. Corresponding images (right) depict OS-17 ERK-KTR reporter C/N localization at initial (T1, red), middle (T2, yellow) and final (T3, blue) timepoints for the indicated cells. Cytoplasmic (red outline) and nuclear (grey outline) masks are shown at T1 for orientation. Scale bar = 25 µm. (F) Representative snapshot maximum intensity projection images of OS-17 reporter cells (left) in the LungSITE model, following 24hr exposure to indicated treatments, and showing relative ERK-KTR (yellow), FIRE (turquoise), and Hoechst staining under these conditions. (G) Quantification of ERK-KTR activity (top), measured as the C/N ratio with normalized nuclear FIRE expression (bottom) at 24hrs post-treatment. Mean of separate experimental replicates are indicated by different symbols, overall mean of experiments is represented by the horizontal line. Data was obtained from 3 independent replicate experiments. Scale bar = 50 µm. (H) Representative snapshot maximum intensity projection images of OS-17 reporter cells extracted from a continuous live-imaging session in LungSITE for 20hrs. Individual images depict changes in ERK-KTR (yellow) localization and variation in FIRE (turquoise) signal intensity. Cells were counterstained with Hoechst (magenta). Scale bar = 50 µm.

We next confirmed the ability of reporter engineered OS-17 lines to colonize murine lungs by lateral tail vein inoculation followed by an incubation period of 2-weeks. (**Figures 1C-D**). Explanted and sectioned lung tissues bore multiple metastatic lesions per section and varying size distributions (1 to >64 cells/cluster), reporters were readily identifiable, and these features were reproducible (**Figures 1C-D**). Within the tumor population over time, observed death events were infrequent under baseline culture conditions, with 23 deaths manually identified among 336 tracked cells during the first 20 hours of imaging. OS-17 cells migrated within their niches and displayed changes in ERK reporter localization status over the course of 7-days (**Supplemental Figure 1A**). Likewise, Calcein AM and MitoSpy staining of the lung revealed maintenance of viability for at least 7-days post-tissue harvest (**Supplemental Figure 1B**). These data confirmed the ability of engineered reporter cells to form orthotopic lung metastasis, consistent with the behavior of unmodified OS-17 cells and maintain viability within the LungSITE model for extended periods.

To segment and track individual reporter cells in LungSITE we trained custom Cellpose models as previously described (30). Using the ratio of C/N fluorescence intensities, reporter activity was calculated and plotted at the single cell level then visually cross-checked with imaging data over time to confirm accuracy (**Figure 1E, Supplemental Figure 1C**). We then performed a qualitative comparison of reporter function in LungSITE (**Figure 1F**) relative to in vitro validation experiments (**Figure 1B**). Under control conditions (growth-factor free media), we observed modest levels of mean ERK reporter activity and heterogeneous FIRE expression within individual tumor cells/nodules (**Figures 1F-G**). These data aligned with prior observations where we have shown that low to moderate ERK activity produces Fra-1 expression heterogeneity because signaling becomes intermittent, pulsatile, and asynchronous from cell-to-cell (6,29,37). Addition of exogenous FGF2 or MEKi revealed expected ERK-KTR and FIRE activation/suppression in tissue (**Figures 1F-G**), consistent with 2D culture (**Figure 1B**). Finally, longitudinal reporter tracking in LungSITE demonstrated evidence of spatial and temporal fluctuations in ERK signaling and FIRE stabilization (**Figure 1H**). Altogether, these data confirmed that LungSITE is capable of accurately tracking and reporting time and space varying ERK and FIRE reporter activity in osteosarcoma lung lesions *ex vivo*.

### Temporal variation in ERK activity and FIRE expression correlate with cell fate following drug challenge with MCL1 inhibitor

Having validated reporter cell function in LungSITE, we next sought to determine the relationship between ERK signaling, FIRE expression, and single cell drug response in the context of the lung microenvironment. Because ERK signaling is a key regulator of cell fate through various mechanisms, including via modulation of prosurvival MCL1 abundance and function (38–40), we hypothesized that spatial and/or temporal variation in ERK signaling may directly influence susceptibility to MCL1 inhibitors (MCL1i). To test this hypothesis, we inoculated NOD.Cg-*Prkdc^scid^*/J mice with OS-17 reporter cells. 2-weeks post-inoculation, lung tissues were harvested, sectioned and live-imaged. Under vehicle control conditions tumor cells persisted with no apparent changes in tumor morphology or cellular abundance (**Figure 2A**, top panels). By contrast, lesions treated with MCL1i (500 nM) underwent significant cell death by 20 hours (**Figure 2A**, bottom panels). However, we observed a fraction of persisting cells, which were observed in 84% of lesions by the end of a 20 hour treatment period, that were delineated by active movement and maintenance of cytoplasmic and nuclear morphology and survived the full imaging time window (**Figure 2A**, see arrows).

**Figure 2.**
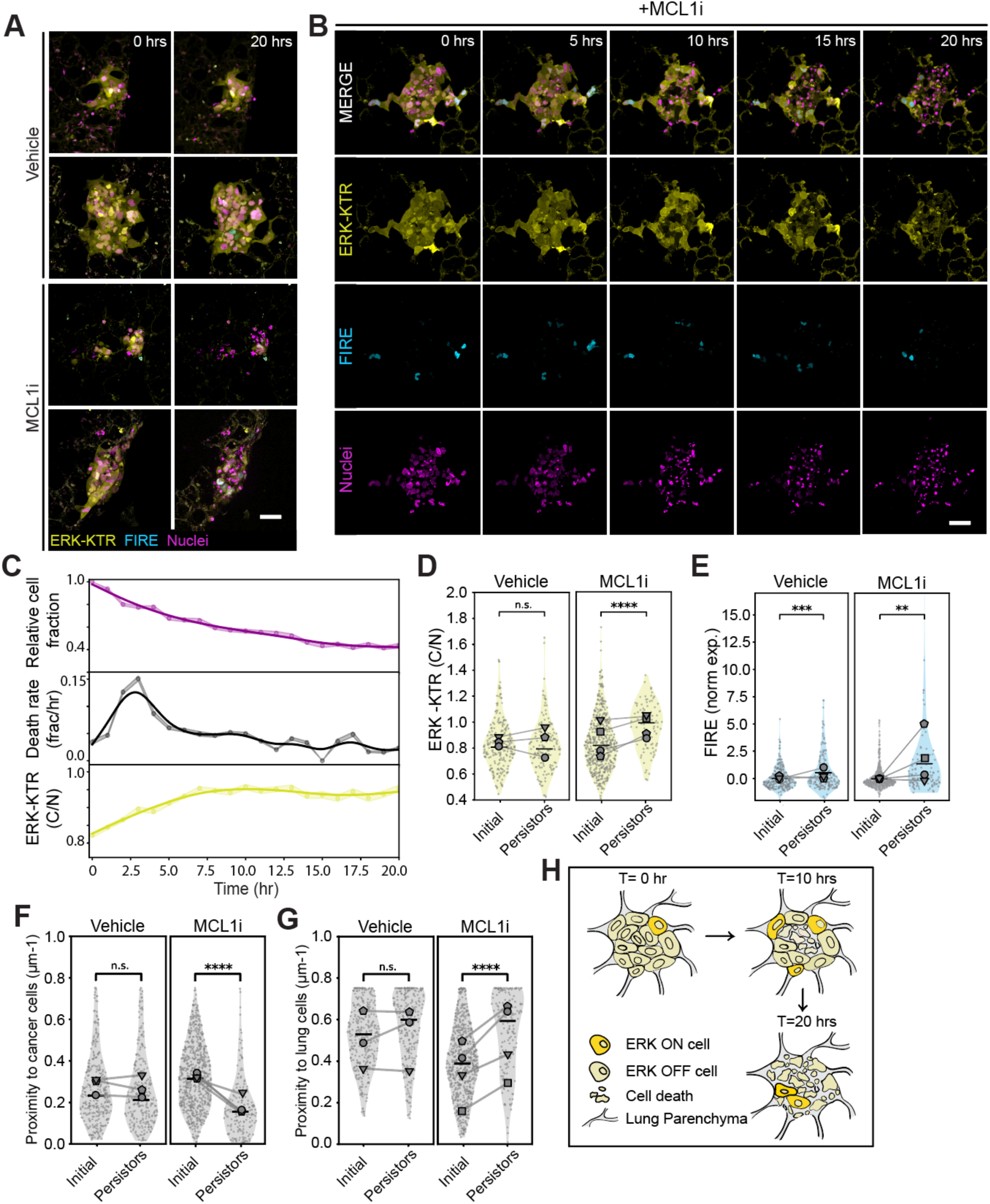
Resistance to MCL1 inhibitor induced cell death correlates with spatiotemporal ERK signaling state. **(A)** Representative snapshot maximum intensity projection images of OS-17 reporter cells live-imaged in LungSITE exposed to Vehicle or 500nM MCL1 inhibitor, AZD5991. Images show ERK-KTR (yellow), FIRE (turquoise) and nuclei, indicated by an H2B reporter (magenta). Scale bar = 50 µm. **(B)** Representative snapshot maximum intensity projection images of OS-17 cells live-imaged in LungSITE treated with 500nM MCL1 inhibitor, showing cell death represented by changes in ERK-KTR (yellow) and FIRE (turquoise) signal intensities and formation of pyknotic nuclei (H2B reporter, magenta). Scale bar = 50 µm. **(C)** Line graphs depicting remaining cell fraction over time (top, magenta line), death rate as fraction per hour (middle, black line), and ERK-KTR activity as a function of C/N ratio (bottom, yellow line). Data was obtained over the indicated time course following treatment with 500nM Mcl1 inhibitor, AZD5991, and is the aggregate of 4 biological replicates. Shaded areas represent the bootstrapped 95% confidence interval (CI). **(D)** Violin plots depicting ERK-KTR activity (C/N ratio) of OS-17 cells in LungSITE at treatment initiation and 20hrs post-treatment. Mean of separate experimental replicates are indicated by grey symbols, the aggregate mean is indicated by the horizontal line. The connecting line between symbols represents the initial and persister activities of each replicate. **(E)** Plots representing normalized FIRE expression, **(F)** relative proximity of an individual living cancer to other cancer cells, or **(G)** relative proximity of an individual living cancer cell to the lung at treatment initiation and 20hrs post-treatment. Relationships were extracted from the dataset presented in panel **(D)**. **(H)** Schematic of death and ERK activation patterns in the lung under MCL1 inhibitor treatment highlighting that ERK activity due to signals received from the lung in dark yellow “ERK ON” cells was heterogeneous. Following treatment, core cells with low ERK activity began dying, followed by edge cells and emergence of a resistant cell population featuring adaptive ERK activation.

Within the surviving tumor cell population, we noted a trend of increased ERK-KTR activity and FIRE expression, and these cells tended to reside closer to the tumor-lung interface (**Figure 2B**). To quantify this relationship, we plotted death rates and ERK-KTR activity over time across multiple experiments revealing a striking temporal correlation between onset of ERK biosensor activity, slowing of cell death rates, and increased survival probability (**Figure 2C**). Quantitative measurement across replicates revealed that, on a single cell basis, ERK-KTR activity and FIRE expression were significantly correlated with survival following MCL1i treatment (**Figures 2D-E**). Furthermore, on average, these cells tended to reside significantly father away from cancer cells and closer to the tumor-lung interface at the end of the 20-hour data collection window (**Figures 2F-G**). To ensure that surviving did not simply die beyond the 20-hour data collection window, we filmed lesions continuously for a period of 7-days. Consistent with our initial findings, the majority of these persister cells remained for the full 7-day period, maintained reporter expression, nuclear morphology, and active membrane dynamics, similar to imaged control cells, despite the presence of inhibitor in the media for the duration of the experiment (**Supplementary Figure 2A**).

Finally, to rule out the possibility that drug tolerant cells simply represent a unique clone with special properties within multiclonal lesions, we engineered OS-17 cells to express histone H2B fused to either mTurquoise, mVenus, mCherry, or iRFP670. Equal proportions of each engineered cell were then co-injected by tail vein and tumor bearing lungs harvested (**Supplementary Figure 2C**). Intra-lesion histone color patterns were then used as a proxy for clonality, revealing that approximately 70% of lesions were monoclonal and likely arose from a single seed, <30% composed of 2 clones, <1% composed of 3 clones, and no 4 clone lesions detected (**Supplementary Figure 2C-D**). By contrast, persister cells were observed in 52% of imaged lesions at 7 days (**Supplementary Figure 2A)**, thus inconsistent with a hypothesis that rare special clones give rise to persister cells. Instead, supporting a conclusion that persister cells arose largely from adaptive resistance mechanisms that contributes to space and time varying single cell death rates.

### Coordinated tumor-tumor and tumor-lung interaction dynamics underly adaptive ERK signaling and death-persistence fate decisions

Our data suggested that both ERK activity state and spatial position may be significant determinates of cell fate contributing to intratumoral drug response heterogeneity. Qualitatively, tumor cells with increased ERK-KTR activity and FIRE expression were often observed closest to the lung and died relatively later, or persisted, compared to cells in the tumor core (**Figure 2B**). To gain a nuanced and quantitative understanding of these relationships, we performed computational analysis and timeseries modeling of the drug response utilizing live-imaging extracted transition probabilities, i.e., Markov modeling. We analyzed trends in tumor-scale ERK-KTR activity relative to individual cell death events on drug (**Figure 3A**), reasoning that in addition to signaling induced by host-derived factors, neighboring cell death events may also influence single cell ERK signaling state. Such findings would be similar to published observations in epithelial sheets and contribute to spatiotemporal tumor cell fate (41). Indeed, we found a time and space dependent ERK-KTR activation relationship in tumor cells adjacent to a death event, spanning ∼80µM and began ∼4 hours after a nearby death event occurs (**Figure 3A**). To extract a more defined relationship between proximal cell death events, ERK biosensor activation, and single tumor cell fate, we generated a ratio metric Sd/R. Sd/R was defined as the sum of the proximities of death events over an 8-hour window to each other surrounding cell, where Sd represents presence of a death event and R represents the distance to the death event in micrometers. Stratifying Sd/R thus captures the ERK biosensor ON (ERK-KTR_LOW_->ERK-KTR_HIGH_) and OFF (ERK-KTR_HIGH_->ERK-KTR_LOW_) state switching rate probability of individual cells as a function of proximity to a cell death event (**Figure 3B**). Cells stratified by moderately elevated Sd/R displayed a multi-fold increase in the likelihood of transitioning from an ERK-KTR_LOW_ to ERK-KTR_HIGH_ state (**Figure 3C**). These data indicated that proximal death events induced an adaptive spatiotemporally-linked ERK response in surrounding cancer cells that superimposed on pre-existing signaling induced by host-derived factors.

**Figure 3.**
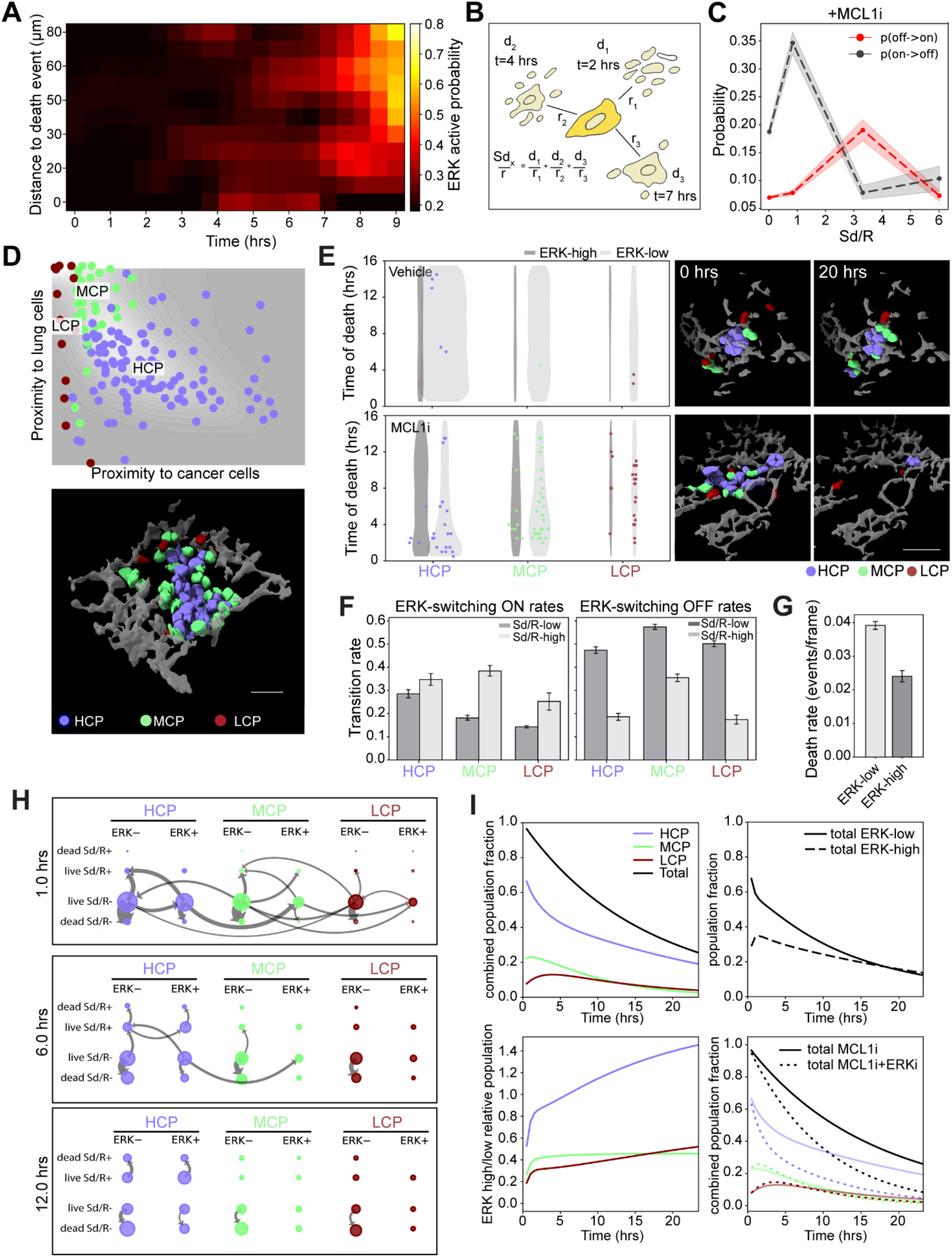
Host and death-associated ERK signaling dynamics underly spatiotemporal cell fate selection. **(A)** ERK activation probability heatmap as a function of distance to a cell death event (µm) and time using data obtained from LungSITE metastatic lesions treated with a 500nM MCL11i. Data are aggregated from 4 biological replicates **(B)** Schematic describing the metric Sd/R, which was used to calculate the probability of death-induced ERK signaling. Sd/R was calculated as the sum of the proximities (inverse distance) of death events over an 8hr window to each cell (see Methods). **(C)** Calculated ERK state probably for activation, off◊on (red) and ERK deactivation, on ◊ off (black) as a function of proximity to death events (Sd/R) in OS-17 LungSITE metastasis treated with 500nM MCLi. **(D)** Scatter plot with a 2D kernel density contour overlay depicts position of cancer cells as a function of proximity to cancer cells and lung cells (top). Cancer cells were accordingly stratified into 3 niche states: high, mixed, and low cancer proximities (HCP in blue, MCP in green, and LCP in red). Example 3-D rendering of a segmented OS-17 metastasis in LungSITE (bottom) with individual cells colored according to contact niche state. Scale bar = 50 µm. **(E)** Violin plots showing time of death of ERK_high_ and ERK_low_ OS-17 cells in LungSITE under Vehicle and 500nM MCL1i conditions, stratified by niche states. Representative images (right) depict 3-D rendered OS-17 metastasis at t=0 and 20hrs after treatments, with individual cells colored according to contact niche states. Scale bar = 50 µm. **(F)** Bar plots showing rates of ERK switching in tumors treated with 500nM MCL1i, stratified by proximity to death events (Sd/R) and contact niche states; high, mixed, and low cancer proximities. **(G)** Bar plot showing average death rates of ERK_low_ and ERK_high_ tumor cells following treatment with 500nM Mcl1i in LungSITE. **(H)** Markov model data demonstrating flux between HCP to MCP and LCP contact niche states 1, 6, and 12 hrs following treatment in LungSITE. (I) Graphs showing experimentally measured changes in contact niche state population fractions over time (top-left), fraction of ERK_high_ and ERK_low_ cells (top-right) over time, and relative ERK_high/low_ populations (bottom-left) over time after MCL1i treatment in LungSITE. The graph bottom-right depicts in silico prediction of changes in fraction of contact niche states over time (dotted lines) under dual inhibition by ERKi and MCL1i superimposed on experimentally collected data (solid lines).

Because we noticed a trend where cell death within lesions often began centrally, then moved by outward procession (**Figure 2B**), we next incorporated cell position and contact information, or “contact niche state”, into our analysis (30). We defined metastable contact niche states via a trajectory modeling approach (42) derived from the distribution around the cell perimeter of the distance to neighboring tumor and host cells for any single tumor cell at a given position in time. These contact niche states captured properties of the local microenvironment around each tumor cell as high, mixed, and low cancer proximities (HCP, MCP, and LCP) (**Figure 3D**). Stratifying tumor populations and recorded death events by contact niche state, and ERK-KTR signaling status, revealed a subpopulation of cells in HCP which were ERK-KTR_HIGH_ and drug resistant over the course of the treatment (**Figure 3E**). Death event proximity increased the ON switching rate, or single-timestep probability, for transitioning from ERK-KTR_low_ to ERK-KTR_high_ and decreased the off-switching rate from ERK-KTR_high_ to ERK-KTR_low_ (**Figure 3F**). When observed cell deaths were integrated over time, we observed that ERK-KTR_low_ cells were about twice as likely to die compared to ERK-KTR_high_ cells overall (**Figure 3G**).

We next constructed a composite state space model where HCP, MCP, and LCP contact niche states were split into ERK-KTR_high_ and ERK-KTR_low_, and live/dead. Switching rates between states in single frames define a transition probability, or a Markov model which defines the frame-to-frame time evolution of the discrete state model. Analyzing the flux between states of the mathematical Markov model over time captured the adaptive response to MCL1i, wherein a flux from HCP to MCP and LCP contact niche states initially occurred along with a rapid flux from ERK-KTR_low_ to ERK-KTR_high_ populations. At 24 hours, the largest subpopulation of surviving cells consisted of ERK-KTR_high_ HCP cells, with flux into this drug-resistant state from Sd/R_low_ ERK-KTR _low_ HCP cells to Sd/R_high_ ERK-KTR_high_ HCP cells in which a proximal death event triggered the transition into ERK-KTR_high_ state (**Figure 3H**). The ratio of ERK-KTR_high_ to ERK-KTR_low_ cells remained mostly stable after the initial hour of the drug response, except in HCP cells, where we modeled a persistent increase. Blocking the transition from ERK_low_ to ERK_high_, in silico, predicted that cytotoxicity in MCP and LCP cells remained similar, but over twice as many HCP cells were sensitive to MCL1i by blocking the adaptive drug response (**Figure 3I**). By resolving contact niche, death event proximity, and ERK reporter signaling status, our data-driven mathematical model isolated the dominant adaptive drug resistance mechanism and pointed to ERK inhibition as a viable strategy in combination with MCL1 inhibition for increased cytotoxicity (**Figure 3I**).

### Drug-induced single cell tumor lysis initiates an adaptive FGFR-ERK-dependent and targetable resistance mechanism

Our data suggested that tumor cell death events can actively shape ERK signaling state and the death/persistence trajectory of adjacent cells. These findings led us to hypothesize that cell death induces or releases bioactive compounds capable of eliciting a local response. To directly test this, we cultured OS-17, OS186, and OS052 lines under standard tissue culture conditions and performed cytokine profiling of their cell lysates in the absence of any drug. We found numerous potential ligands, such as IGF2, FGF2, AREG and others that varied in relative abundance between lines (**Figure 4A**), and are known to activate ERK signaling (6). However, OS-17 (**Figure 1B**), OS186, and OS052 cells exhibited low or no-detectable ERK activity at baseline when cultured (**Figure 4B**) yet displayed significant upregulation of Fra-1 in the surviving fraction post-MCL1i treatment (**Figure 4C**). These data implied that ERK activation in persisting cells must be induced or released during drug-induced cell death to initiate a signaling response and drive downstream Fra-1 expression.

**Figure 4.**
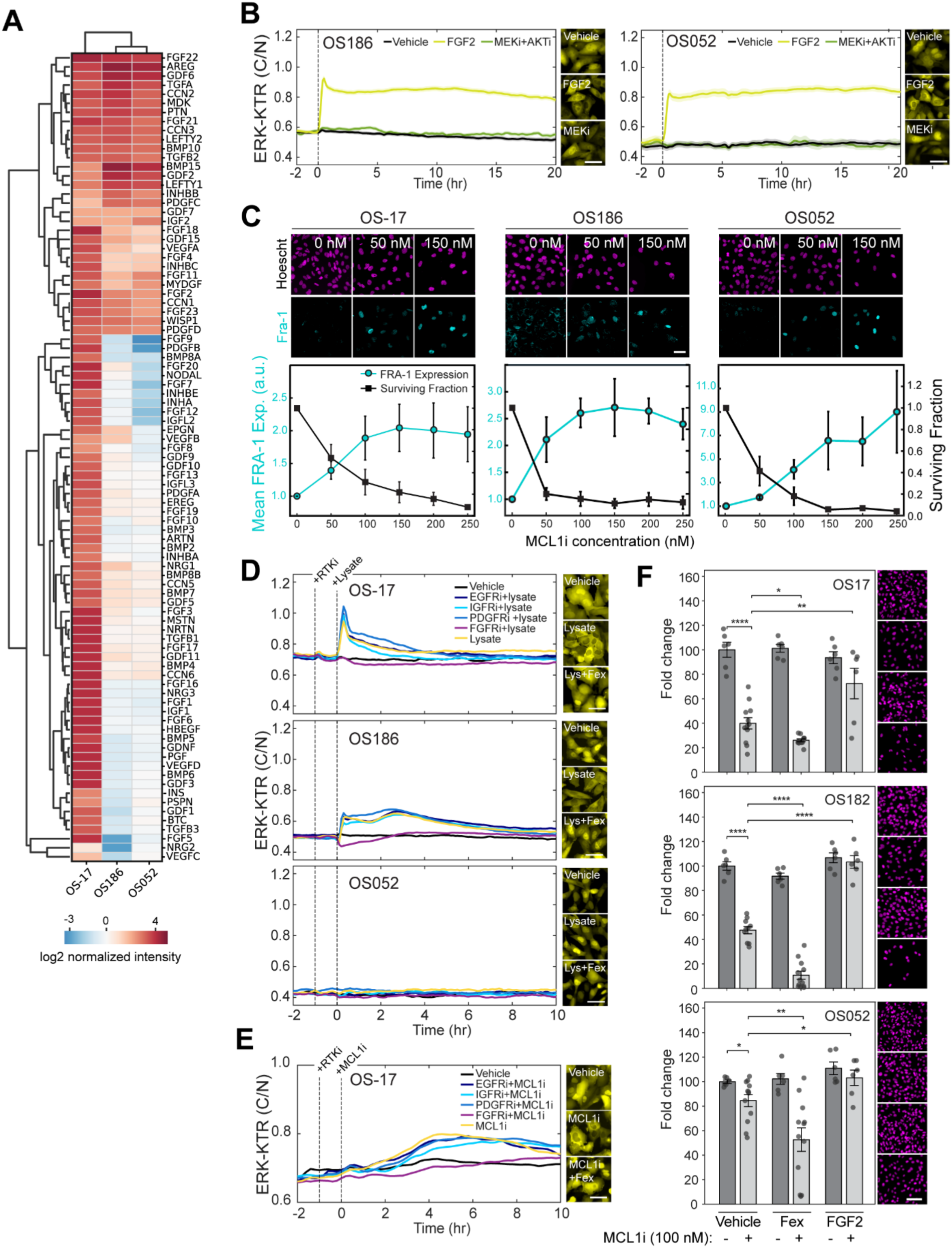
Tumor cell lysis promotes adaptive ERK activation and acquisition of drug resistance. **(A)** Heat map depicting relative secreted growth factor abundance of OS-17, OS052 and OS186 cells measured by cytokine array. **(B)** Mean traces of ERK-KTR activity (C/N ratio) obtained from 2D monocultured of OS186 and OS052 cells over time (solid line), along with inter-quartile range for the specified conditions. Two independent replicate experiments were performed, ≥1500 cells per condition per experiment. Scale bar = 50 µm. **(C)** Mean surviving cell fraction and average normalized Fra-1 expression in OS-17, OS186 and OS052 cells following exposure to varying doses of MCL1i for 24hrs. Images (top) show representative Fra-1 staining (turquoise) for the indicated drug concentrations at 24hrs. Cells were counter-stained with Hoechst (magenta) to delineate nuclei. Data represent the aggregate of four replicate experiments. Scale bar = 50 µm. **(D)** Mean traces of ERK-KTR activity in 2D monocultured OS-17, OS186 and OS052 cells over time exposed to respective tumor cell lysate in the presence of absence of the indicated RTK inhibitors. Vertical dotted lines indicate time of addition for RTK inhibitor (RTKi) or cell lysate. Two independent replicate experiments were performed, ≥2500 cells per condition per experiment. Scale bar = 50 µm. **(E)** Mean traces of ERK-KTR activity obtained from 2D monocultured OS-17 cells treated with the specified RTK inhibitors, followed by 500nM MCL1i.Two independent replicate experiments were performed, ≥6000 cells per condition per experiment. Scale bar = 50 µm. **(F)** Bar plots displaying mean survival fraction of OS-17, OS186 and OS052 cells following 24hrs of treatment with FGF2 or FGFR inhibitor (fexagratinib) alone, or in combination with 100nM MCL1i. Representative images (right) show Hoechst-stained nuclei of remaining cells following 24 hrs of the indicated treatments. Two independent replicate experiments were performed. Scale bar = 50 µm.

To test the ability of cell lysis to induce protective ERK signaling in adjacent cells, we first generated cell lysates from OS-17, OS186, and OS052 cells by mechanical shearing. This approach was used to avoid potential confounding factors caused by adding a drug to induce lysis. Cell line specific lysates were then applied to corresponding ERK-KTR expressing cells in standard monoculture conditions, in the presence or absence of RTK inhibitors targeting FGFR (AZD4547, fexagratinib), EGFR (erlotinib), IGFR (NVP-AEW541), or PDGFR (CP-673451). We observed an immediate and robust ERK response in OS-17 and OS186 cells that was blocked by FGFR but not EGFR, IGFR, or PDGFR inhibitor (**Figure 4D**). By contrast, addition of lysate to OS052 cells did not produce an ERK signaling response, nor did addition of inhibitor significantly suppress of baseline signaling (**Figure 4D**). This finding was unexpected as MCL1i induced Fra-1 expression in prior experiments (**Figure 4C**), suggesting that Fra-1 expression may be driven through ERK-independent mechanisms in some cell types/ patient models (43). Using ERK-KTR expressing OS-17 lines, we then directly treated cells with MCL1i +/-RTK inhibitors and tracked ERK biosensor activity over time. Under these conditions, we found that ERK activity began to spike as cells began to die at ∼2 hours post-MCL1 inhibitor addition, temporally consistent with onset of cell death in our prior LungSITE experiments, and this effect was blocked by FGFR inhibitor (**Figure 4E**).

To solidify that activation of the FGF-FGFR signaling axis functions as an adaptive cell survival mechanism, we tested the ability of FGFR blockade or activation to enhance or protect against MCL1i-induced cell death, respectively. In OS-17, OS186, and OS052 cells, we found that FGFR blockade with 2uM fexagratinib (AZD4547) did not have a significant effect on cell survival alone (**Figure 4F**). However, when combined with the approximate IC50 dose of MCL1i, we observed an increase in cell death in OS-17 and OS186 cells relative to MCL1 inhibitor alone. While a similar increase in cell death was observed for OS052 cells, response between replicates was more heterogeneous and this line was relatively less sensitive to MCL1i alone as compared to OS-17 and OS186 cells. Finally, addition of exogenous FGF2 produced a strong protective effect in all cell lines tested (**Figure 4F**). Additionally, these data showed that while OS052 cells did not display evidence of “self-activating” ERK signaling, FGF-FGFR mediated activation of ERK signaling induced protection in this model.

Finally, we assessed the synergy-additivity relationship of combinatorial FGFRi and MCL1i OS-17, OS186, OS052, and 143B cells. We included 143B cells as a “resistant control” reasoning they would not exhibit synergy or additivity with combinatorial FGFRi-MCL1i because of persistent Ras mutant-driven ERK activity. Consistent with our prior findings (**Figure 4F**), combinatorial treatment produced a synergistic response in OS-17 and OS186 cells, showing Loewe synergy scores of 15.84 and 22.03, respectively, whereas OS052 cells were responsive to MCL1i but insensitive to combination with FGFRi and showed a low synergy score of 0.27 (**Supplementary Figure 3B**). 143B cells were highly insensitive to both FGFRi and MCL1i consistent with a mechanism whereby persistent intrinsic, or induced, ERK signaling promotes cell survival.

### Combinatorial RTK-MCL1 inhibition strategies enhance MCL1i efficacy *ex vivo*

Based on our mathematical model (**Figure 3I**) and drug synergy data (**Supplementary Figure 3B**), we posited that combinatorial FGFR and MCL1 targeting drugs might eliminate an increased fraction of tumor cells in the lung and suppress ERK signaling-mediated adaptive resistance. To this end, we tested inhibitors that ranged in FGFR selectivity, from very selective drugs fexagratinib and erdafitinib, to broad inhibitors such as nintedanib, dovitinib, and pazopanib that co-target FGFR, VEGFR, and PDGFR (**Figure 5A**). Patient populations are likely to have variable sensitivity to RTK-directed therapies underlying our logic to include broad specificity inhibitors. Furthermore, pazopanib and other broad specificity RTK inhibitors have been shown to exhibit efficacy against metastatic osteosarcoma in clinical trials (14–16). We found that each RTK inhibitor alone had minimal impact on tumor cell survival (**Figure 5A-B**). However, co-treatment with 100nM MCL1i, a five-fold reduction in concentration relative to our preceding experiments, resulted in significant cell death compared to single agent RTK or MCL1 inhibitor. We then further evaluated fexagratinib and pazopanib as respective specific- and broad-targeting agents in LungSITE (**Figure 5C**) tracking cell fates over time. Control, 100nM MCL1i, or 2uM RTKi alone displayed minimal to no detectable cell death. However, combinatorial treatment with fexagratinib and MCL1i produced significant cell death (**Figure 5C-D**). Combinatorial efficacy was further accentuated for pazopanib dosed with MCL1i (**Figure 5C-D**), allowing us to kill almost double the tumor cells compared to MCl1i treatment alone. To confirm that receptor-level blockade inhibits adaptive ERK signaling responses in the presence of MCL1i, we monitored single cell ERK-KTR activity following drug exposure using LungSITE. As predicted, fexagratinib and pazopanib inhibited MCL1i– induced ERK activation over time, independent of contact niche state (**Figure 5E, Supplementary Figure 2F**), and ERK-KTR_high_ cells were no longer observed near death events following MCL1i exposure, indicating that combinatorial treatment could block the adaptive response (**Figure 5F**). We then tested the effectiveness of this combination in 143B cells, predicting relative insensitivity due to persistent intrinsic Ras mutant-driven ERK signaling as opposed to adaptive induction. As predicted, low dose MCL1 inhibition had minimal impact on 143B cell survival, alone or in combination with pazopanib (**Supplementary figure 6A, B**). However, direct inhibition of ERK signaling through MEK blockade resulted in a significant decrease in cell survival that was enhanced by addition of low or high doses of MCL1i (**Supplementary figure 6**). Altogether, these data suggest that blockade of adaptive (or persistent) ERK activation is a tool to increase the efficacy of MCL1i while reducing the required drug dose in experimental models.

**Figure 5.**
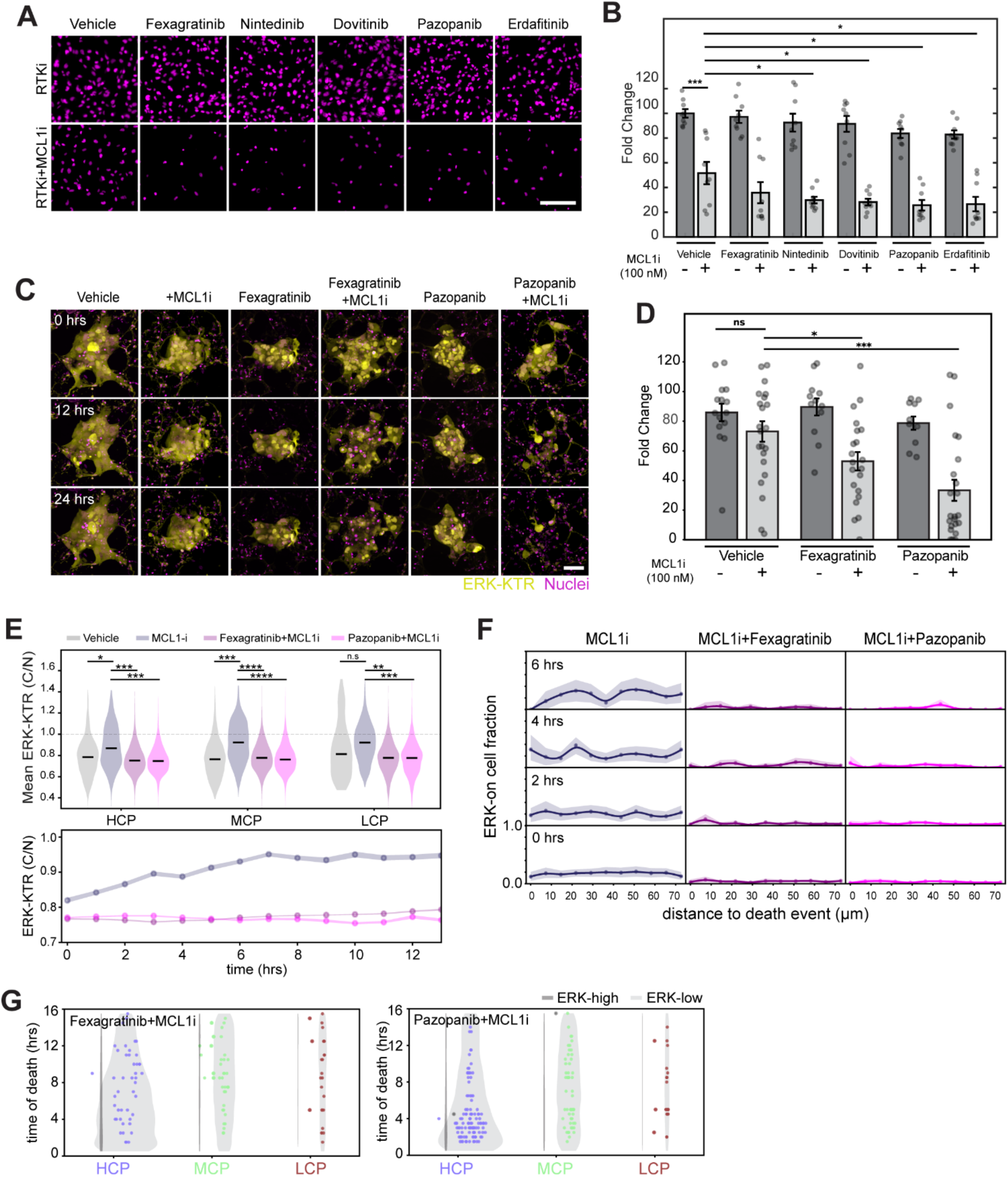
Combinatorial receptor tyrosine kinase and MCL1 inhibitors suppress adaptive ERK-mediated resistance. **(A)** Representative images of Hoechst-stained nuclei of OS-17 monocultured 24hrs following treatment with the indicated receptor tyrosine kinase inhibitors (2µM concentration) alone (top row) or in combination with 100nM MCL1i (bottom row). Scale bar = 50 µm. **(B)** Bar plot showing fold change in OS-17 abundance in monoculture following 24 hrs of treatment with receptor tyrosine kinase inhibitor alone or in combination with 100nM MCL1i. Error bars present the Standard error of mean, while individual dots present technical replicates. Data represents the aggregate of 3 biological replicates. **(C)** Representative maximum intensity projection images of OS-17 cells, ERK-KTR (yellow) and nuclei (magenta), indicated by H2B reporter and Hoechst stain, in LungSITE model treated with 100nM MCL1i alone and in combination with 2µM fexagratinib or pazopanib, are shown at different timepoints. Scale bar = 50 µm. **(D)** Bar plot showing fold-change in cell count of OS-17 cells in LungSITE following 24hrs exposure to the indicated drug concentrations. Data was aggregated from 3 independent replicate experiments. Error bars present standard error of mean, while dots present fold change of individual lesions **(E)** Violin Plot (Top) showing distribution of mean ERK-KTR activity (C/N ratio) of tumor cells in LungSITE following exposure to Vehicle, 100nM MCL1i, or 2µM fexagratinib or pazopanib in combination with 100nM MCL1i stratified by contact niche states. Line plot (bottom) comparing ERK-KTR activity the indicated time-period following treatment with 500nM MCL1i, or 100nM MCL1i in combination with 2µM fexagratinib and pazopanib. Shaded areas represent the bootstrapped 95% confidence interval (CI). **(F)** Line plots showing mean fraction of “ERK ON” cells at different time points as a function of distance from neighboring death events following treatment with 500nM MCL1i, or 100nM MCL1i in combination with 2µM fexagratinib and pazopanib. Shaded areas represent the bootstrapped 95% confidence interval (CI). **(G)** Violin plots of time of death for ERK_high_ and ERK_low_ OS-17 cells in LungSITE in response to 2µM fexagratinib or pazopanib in combination with 100nM MCL1i and stratified by niche states: high, mixed, and low cancer proximities.

### Combinatorial RTK-MCL1 inhibition strategies enhance drug response in vivo

To determine if combinatorial RTK-MCL1 inhibition could improve therapeutic efficacy of MCL1i *in vivo*, we injected mice with luciferase expressing OS-17 cells (OS-17 FLUC) via tail vein and allowed experimental metastasis to form. Three weeks post injection, the mice were randomized based on baseline luminescence into four treatments groups: vehicle, MCL1i, pazopanib and a combination of pazopanib and MCL1i and placed on treatment for 2-weeks. Tumor growth was assessed as the log2 fold change in luminescence relative to treatment start. Vehicle and pazopanib treated mice showed a marked increase in luminescence, indicating substantial tumor growth (**Figure 6A-B**). MCL1i-treated mice showed attenuated tumor growth compared to the vehicle- and pazopanib-treated groups, yet these differences did not reach statistical significance. Combination treatment group showed a decreased fold change to starting luciferase activity, suggesting tumor regression. This change was statistically significantly compared to both the vehicle- and pazopanib-treated groups. When analyzed on the level of individual mice, both MCL1i and combination-treated mice showed similar decrease in luminescence (**Figure 6C**). However, by the end of second week, 80% of the MCL1i treated mice displayed a measurable increase in luminescence compared to only 40% in the combinatorial arms. Although the difference between MCL1i and combinatorial treatment arms were not found to be statistically significant at endpoint (2-weeks) these data are consistent with our *ex vivo* data (**Supplementary Figure 7**) suggesting that combinatorial treatment eliminates a small but persisting fraction of tumor cells that fail to be eliminated by MCL1i alone.

**Figure 6.**
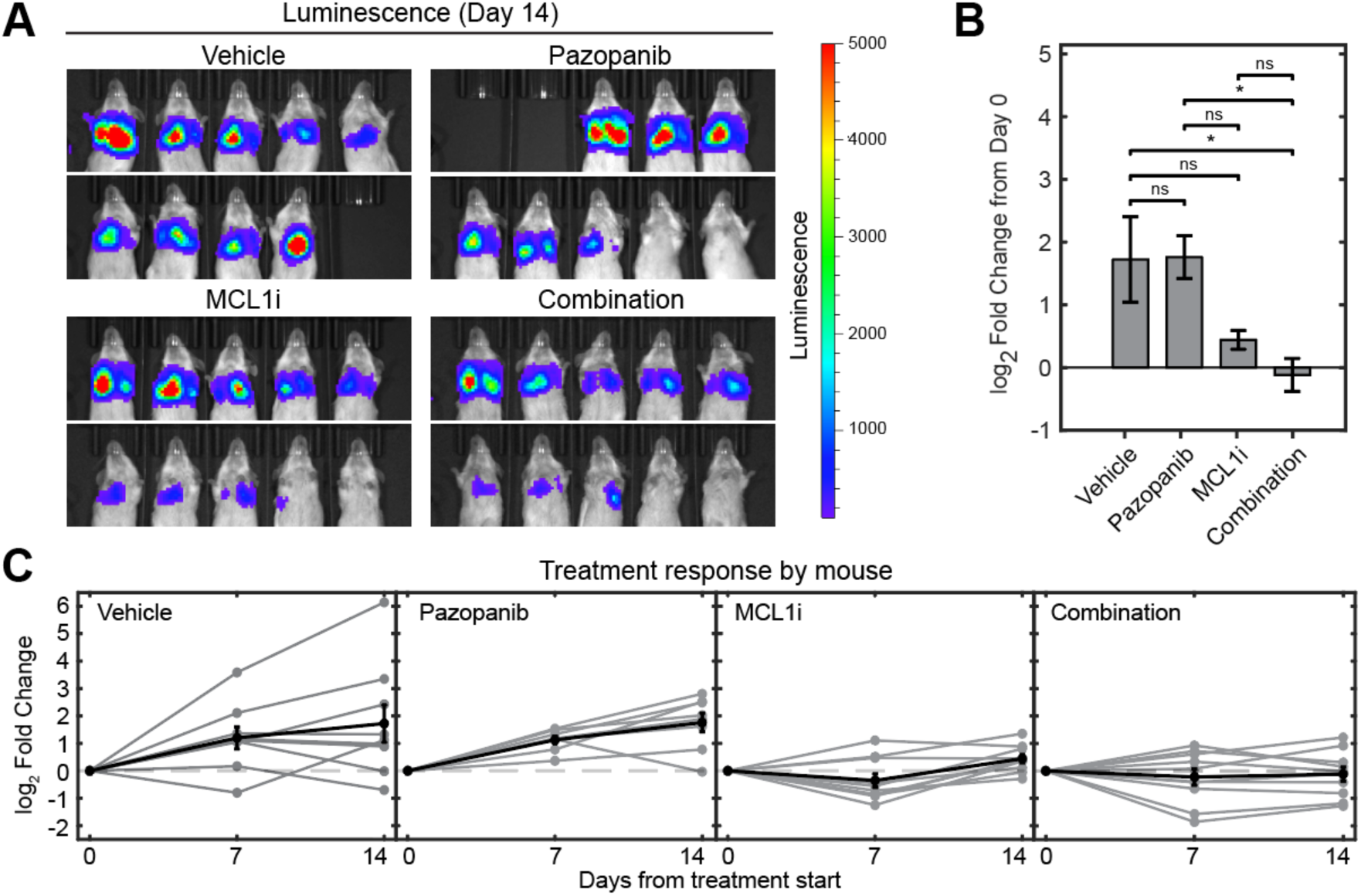
In vivo efficacy of combinatorial pazopanib and MCL1 inhibitor. **(A)** Luminescence imaging of OS-17-LUC tumors in the indicated treatment arms 14 days after treatment initiation. All images were scaled to the same maximum and minimum luminescence. **(B)** Bar chart showing mean log2 fold change in luminescence at day 14 compared to day 0 of treatment administration. Error bars represent standard error of the mean. **(C)** Log2 fold-change in luminescence over the treatment course of individual mice (grey lines) within each treatment. Black lines represent mean log2 fold-change. Error bars represent standard error of the mean.

### FGFR signaling or drug exposure induced upregulation of common drug resistance pathways and correlated with expression patterns in heavily pretreated patient samples

We sought to understand how adaptive ERK activation could promote resistance through downstream effectors. To this end, we performed mass spectrometry on OS-17 cells treated with MCL1i, FGF2, or MEKi in monoculture conditions. MCL1i and FGF2 treated samples displayed strikingly similar patterns of target upregulation, with the most significant changes in Fra-1, CCN2, ERRFI1, ENPP1, and HMOX1 (**Figure 7A**). Interestingly, ENPP1 is a regulator of soft tissue mineralization, confers local immune suppression in cancer, and is associated with aggressive disease and poor prognosis in many solid tumors (44,45). HMOX1 is a bona fide resistance gene regulated by the transcription factor Nrf2, activation of which is correlated with widespread drug resistance across cancer types and a recently described target in osteosarcoma (46). To validate our mass spectrometry data, we performed confirmatory immunofluorescent staining of Nrf2 and HMOX1, revealing that FGF2 or MCL1i treatment resulted in significant induction of both proteins (**Figure 7B**). In 143B cells that exhibited basal resistance to MCL1 inhibitors and persistent ERK hyperactivation, Nrf2 and HMOX1 levels were unaffected by FGF2 or MCL1i, but significantly reduced by MEKi (**Figure 7C**), consistent with high basal expression.

**Figure 7.**
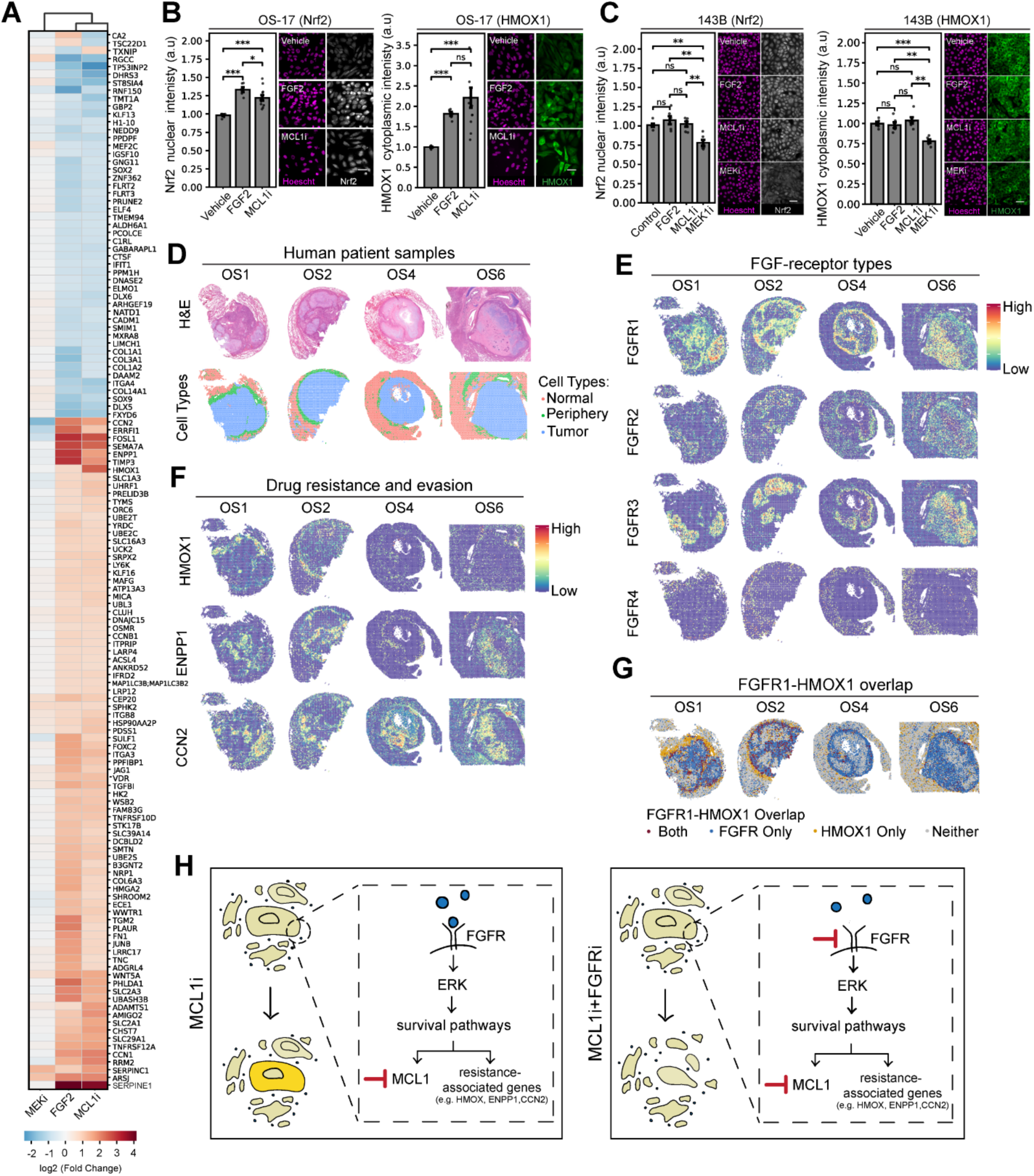
Adaptive FGFR-driven ERK signaling correlates with upregulation of pan-drug resistance genes. **(A)** Heatmap of protein differential abundance in OS-17 cells treated with MEKi, FGF2, or 500nM MCL1i for 24 hours. Scale bar represents Log2 fold-change. Data was obtained from 3 biological replicates. **(B)** Bar plots report normalized nuclear Nrf2 expression (left) and normalized cytoplasmic HMOX1 expression (right) in OS-17 cells in response to Vehicle, FGF2, or 100nM MCL1i for 24 hours. Representative images depict Nrf2 and HMOX1 expression in cells, which were counter-stained with Hoechst (magenta). Data was obtained from 2 biological replicates. Scale bar = 50 µm. **(C)** As in panel (B) Nrf2 and HMOX1 expression are reported for 143B cells in response to vehicle, FGF2, MCL1i or MEKi and representative images included for visualization. Data was obtained from 2 biological replicates. Scale bar = 50 µm. **(D)** H&E-stained 10X Visium spatial transcriptomics osteosarcoma samples and spatial map of the associated cell type annotation for each spot defining tumor (blue), tumor periphery (green), and surrounding normal lung tissue (orange). **(E)** Spatial expression of FGF-receptors and **(F)** drug resistance and evasion associated genes in each of previously introduced samples. **(G)** FGFR1-HMOX1 overlap mapping as a function of median expression (see Methods). Regions with dual expression above median are indicated in dark red, FGFR1 only in blue, HMOX1 only in dark yellow, and no detectable expression of either in gray. **(H)** Schematic of proposed FGFR-mediated adaptive ERK resistance mechanisms in osteosarcoma lung metastasis.

Finally, we queried existing spatial transcriptomics data from heavily pretreated patient lung metastasis (**Figure 7D**) for evidence of resistance pathway expression patterns (7,47). Strikingly, we found that FGF receptors are significantly upregulated in tumor samples, often with distinct spatial patterning by receptor type (**Figure 7E**), suggesting the presence of the necessary receptors for adaptive FGFR-ERK signaling-mediated resistance to occur. FOSL1 expression was poorly captured in these datasets (data not shown). However, profiling of other mass spectrometry identified targets other known ERK responsive genes, CCN2, ENPP1, and HMOX1 levels also revealed significant intratumoral upregulation in patient samples, and often visually overlapped with FGFR expression (**Figure 7E**). Because these samples were heavily pretreated at the time of collection, we could not discern adaptive/persistent versus pre-existing expression of these targets (see Discussion). Nevertheless, FGFR1 and HMOX1 were visibly enriched at the tumor edge, where we overserved adaptive resistance in LungSITE experiments. Using a quantitative overlap algorithm, we found a high degree of overlap between FGFR1 and HMOX1 (**Figure 7G**), although HMOX1 expression varied from sample to sample, and with HMOX1 expression extending into the periphery into the tumor-lung interface, a region known to be subject to high oxidative stress loads (48,49). Altogether, these data were consistent with a mechanism of drug-induced adaptive resistance via dynamic activation of FGFR-ERK signaling and downstream drug resistance pathways following MCL1 inhibitor exposure or clinical treatment regimens (**Figure 7H**).

## DISCUSSION

Here we applied *ex vivo* live-cell imaging and biosensor approaches, using LungSITE (30), to directly observe single cell drug response heterogeneity within living osteosarcoma lung metastases, revealing an adaptive ERK-mediated resistance mechanism. Tracking drug response in real time showed that MCL1 inhibition produced extensive cell death yet left a resistant population linked to ERK cell signaling state, tissue position, and nearby death events. Importantly, these data provide evidence that phenotypic drug response heterogeneity is deterministic rather than stochastic and therefore both discoverable and targetable.

The data presented here showed that RTK-driven environmental signals and drug-induced stress can converge on shared resistance-associated states in metastatic osteosarcoma. We previously showed that lung-derived growth factors activated RTK-ERK regulated survival programs, culminating in upregulation of MCL1, which acted to promote osteosarcoma engraftment and colonization in the lung (6). Here, we found that treatment-induced cell lysis reinforced activation of this same circuitry, amplifying the host response via FGFR-Ras-ERK, which resulted in enhanced tumor cell survival under therapeutic stress conditions. Direct induction of FGFR-ERK signaling via FGF2 exposure or addition of MCL1 inhibitor produced strikingly similar protein responses, including induction of FOSL1/Fra-1, CCN2, ERRFI1, ENPP1, and HMOX1 together with NRF2. CCN2 and ENPP1 are both known regulators of bone development and mineralization, indicating that signaling or therapeutic stress could activate programs associated with normal development (44,50,51), and in the case of ENPP, a mechanism of immune evasion in the cancer context (45). In patient metastases, spatially associated FGFR1 and HMOX1 expression was evident, which adds clinical relevance to our findings. However, patient specimens cannot establish when these programs arose and will require additional functional studies to unravel. Additionally, the translatability of MCL1 inhibitor-induced adaptive resistance mechanisms to other chemotherapeutic or targeted therapies remains to be determined; further studies will be required to systematically understand how other existing and investigational osteosarcoma treatments similarly, or differentially, induce adaptive resistance.

Therapeutically, our data support pairing cytotoxic, or ‘targeted’ cytotoxic agents, with inhibition of the adaptive signals that protect surviving cells. Recent studies of MCL1 inhibitors in preclinical osteosarcoma models support this conclusion, and RTK efficacy has been demonstrated in clinical trials (14,17,19,22,38). In our models, selective and broad RTK inhibitors had little activity alone yet robustly suppressed MCL1 inhibitor-induced ERK activation and increased tumor killing even at reduced MCL1 inhibitor concentrations. Dose-sparing strategies are attractive given the toxicity of many existing and experimental therapies, although improved tolerability must be established directly as RTK inhibitor combinations may introduce new toxicity risks.

In this study, we specifically investigated MCL1 inhibition as a target of interest downstream of microenvironment-driven survival signaling based on our prior findings (6). Going forward our results justify additional and expanded studies, spanning multiple RTK-driven pathways in combination with different targeted or cytotoxic drugs, to define broadly efficacious and patient-specific drug combinations that suppress residual-cell survival and provide sustained disease control. Identification of patient-specific combinations will be essential, as we found some level of variation in adaptive resistance within the relatively small sample size of patient-derived cell line models tested here. Despite such variation, we demonstrated the larger principle that identification and interception of treatment-induced adaptive signaling can improve the depth, durability, and therapeutic index of drug response.

A limitation of the study presented here is lack of a complex immune system *ex vivo* and *in vivo*. Immune cells provide rich and dynamic sources of microenvironmental signaling, particularly macrophage populations, which are often highly enriched in osteosarcoma lung metastasis (7). Future studies will require the integration of living immune-competent tissues and patient-samples to uncover the contributions, and precise modes, of immune-tumor signaling interactions contributing to adaptive resistance. Such studies are made possible using the experimental and computational approaches developed and applied herein.

Finally, the mechanistic insights presented here were made possible by live-cell tracking in tissues coupled to data-driven state-space modeling which enabled precise connection of tumor cell position and neighboring death events to ERK signaling state transitions and cell fate. This was accomplished by converting live 3-dimensional movies into interpretable data sets via cell identification and quantification in dense and complex tissue environments, which despite the strides made here, remains a substantial ongoing challenge in the field of image analysis. Still, extraction of mechanistic relationships from our dynamic data highlights a major advance enabled by the LungSITE platform. Continued advances in computer vision and lineage reconstruction should improve the scale and precision of these analyses (52,53). Such advances will contribute to a growing body of literature that continues to recontextualize the metastatic osteosarcoma microenvironment as a dynamic entity; revealing increasingly nuanced mechanic details that will inform a previously inaccessible biological understanding of osteosarcoma and reveal new translational opportunities.

## ACKNOWLEDGEMENTS

We would like to thank Dr. Troy McEachron for providing access to patient spatial transcriptomics datasets. The research reported in this publication was supported by funding support to RM and AED from the Knight Cancer Institute and to AED from National Institutes of Health, Office of Research Infrastructure Programs, under award number K01OD031811; the National Cancer Institute, under award number 1R01CA300732; Sam Day Foundation; MIB Agents/Fishin’ For The Cure OutSmarting Osteosarcoma. The research reported in this publication used computational infrastructure supported by the Office of Research Infrastructure Programs, Office of the Director, of the National Institutes of Health under Award Number S10OD034224. The content is solely the responsibility of the authors and does not necessarily represent the official views of the National Institutes of Health. We acknowledge assistance by staff in the Advanced Multiscale Microscopy Shared Resource, supported by the OHSU Knight Cancer Institute NIH P30 CA069533 and the OHSU University Shared Resources Program. Equipment purchases included support from the OHSU Foundation, the Knight Cancer Institute, the MJ Murdock Charitable Trust, and the Collins Medical Trust. Artificial intelligence was used for editing this manuscript. All data, content, and conclusions were generated, interpreted, and described solely by the listed authors.

## EXTENDED MATERIALS AND METHODS

### Cell culture

OS-17 cells were derived from an early passage of the OS-17 patient-derived xenograft obtained from a primary femur biopsy performed at St. Jude’s Children’s Research Hospital in Memphis and grown in RPMI (Gibco, 11875093) supplemented with 10% FBS (31). OS052 and OS186 were obtained from the Sweet-Cordero lab in UCSF and grown in DMEM (Gibco, 11995-065) supplemented with 10% bovine growth serum (Cytiva, SH30541.03) (54). 143B cell line was purchased from ATCC and grown in DMEM supplemented with 10% FBS.

### Reporter line generation

OS17 reporter cell line was generated through lentiviral transduction of ERK translocation reporter (ERK-KTR) fused to Clover (33), Fra-1-based Integrative Reporter of ERK (FIRE) fused to mTurquoise (27) and histone H2B fused to iRFP670 cloned into the pLJM1 lentiviral backbone (34). Cells were then selected for reporter integration using 1 µg/mL puromycin (Gibco, A1113803) and sorted by flow cytometry. The 143B, OS052 and OS186 reporter cell lines were generated by lentiviral transduction of ERKTR-iRFP (55) and H2B-mCherry2 in the pLJM1 backbone followed by selection using puromycin (1 µg/mL, Gibco, A11138-03). OS-17-FLUC cells (6) were generated through lentiviral transduction of Firefly luciferase (Cellomics Technology, PLV-10064-50). OS-17-H2B reporter cells were generated by lentiviral transduction of H2B fused to mVenus, iRP670, mTurquoise or mCherry in the pLJM1 backbone followed by selection using geneticin (200 µg/mL, Gibco, 10131035).

### Experimental metastasis and generation of precision cut lung slices

Cell suspensions of 1 × 10^6^ OS17 or 143B reporter cells were injected via tail-vein into anaesthetized 8–10-week NOD.Cg-*Prkdc^scid^*/J mice. For H2B clonality experiments, 1.5 × 10^5^ of OS17-H2B mVenus, iRP670, mTurquoise and mCherry were mixed into a single cell suspension and used for injection(1 × 10^6^ total). Mice were humanely euthanized 2 to 3 weeks post injection. Lungs were inflated within the chest cavity with 2.5% low point melting temperature agarose (Thermo Scientific, R0801) prepared with DMEM:F12 phenol-free media (Gibco, 11039047). Lungs were removed and kept on ice until slicing. Lungs were sliced at 300 µm thickness using Compresstome VF-510-0Z in ice cold advanced DMEM/F-12 (Gibco, 12634010). Lungs were then cultured in DMEM/F-12 supplemented with insulin (250 ng/ml, Sigma-Aldrich, I1882), transferrin (10 µg/ml, Sigma-Aldrich, T8158), β-estradiol (10^-10^ M, Sigma-Aldrich, E2758), hydrocortisol (1.4 × 10^-6^ M, Sigma-Aldrich, H4001), sodium selenite (2.6 ng/ml, Sigma-Aldrich, S5261), vitamin C (0.1 µg/mL, Sigma-Aldrich, A4403), vitamin A (0.2 mg/mL, Sigma-Aldrich, R2625) and primocin (100 µg/ml, InvivoGen, Ant-pm-1). Media was changed every other day.

### Live-cell microscopy

All Imaging was done using Nikon Ti2 automated microscope fitted with an Oko-Lab environmental chamber maintained at 37 °C and 5% CO_2_, SOLA II LED illumination source, Orca Fusion BT camera, and controlled with NIS-Elements AR software.

For imaging cell line monocultures, glass-bottomed 96 well plates (Cellvis, P96-1.5H-N) were coated with rat tail collagen (Corning, 354236), then washed with PBS (Gibco, 14190144). Cells were seeded at a density of 7,000 cells per well and incubated overnight in complete growth medium. Cells were then washed twice with phenol-free DMEM/F12 and then placed in imaging media consisting of phenol-free DMEM/F12 supplemented with transferrin (10 µg/ml), hydrocortisol (1.4 × 10^-6^ M), β-estradiol (10^-10^ M) and Hoechst (Thermo Scientific, 62249, 1:50,000). Plate was left to incubate for 4 hours before imaging. Images were taken every 6 minutes. MATLAB was used to analyze generated data.

For imaging PCLS, PCLS were transferred to a #1.5 glass bottomed 24 well plate (Cellvis, P24-1.5H-N) and clamped down to the bottom of the plate with custom laser cut acrylic clamps to flatten the tissue and prevent its movement during imaging. PCLS were washed twice with phenol-free DMEM/F12, for 10 minutes per wash, then placed in PCLS imaging media consisting of phenol-free DMEM/F12 supplemented with transferrin, hydrocortisol, β-estradiol, vitamin C, vitamin A and primocin. Hoechst (Thermo scientific, 62249) was added to the imaging media to visualize lung cell nuclei and to ensure nuclear identification in lesions with variable H2B reporter expression. Inhibitor and growth factor treatments were added around 6 hours prior to imaging to allow their full effects to take place before imaging, except for Mcl1 inhibitor AZD5991, which was added directly at the start of imaging. Images were taken every 20 or 30 minutes for a duration of 20 – 48 hours, depending on the experiment set. For experiments longer than 48 hours, imaging media and included treatments were replenished every other day. SITE analysis pipeline was used to analyze generated data. Replicate experiments used PCLS obtained from different mice.

Treatments used included FGF2 (100 ng/ml), PD0325901 (100 nM, Selleckchem, S1036), MK-2206 (1 µM, Selleckchem, S1078), AZD5991 (Selleckchem, S8643), fexagratinib (2 µM, Selleckchem, S2801), NVP-AEW541 (2 µM, Selleckchem, S1034), erlotinib (2 µM, SelleckChem, S7786), CP-673451 (2 µM, Selleckchem, S1536), pazopanib (2 µM, MedChemExpress, HY-10208), Dovitinib (2 µM, MedChemExpress, HY-50905), erdafitinib (2 µM, MedChemExpress, HY-18708), nintedanib (2 µM, MedChemExpress, HY-50904) and cell line derived lysates (10 µg per well, methods below). To determine the concentrations of the MCL1 inhibitor (AZD5991) used in subsequent experiments, dose– response titration assays were performed on OS-17 cells, and the IC50 for cell viability was determined. The IC50 concentration was selected as the lower dose (100 nM), while a five-fold higher concentration was used as the high-dose condition (500 nM). To account for potential variability between individual MCL1i vials, dose–response titrations were performed independently for each vial, and the corresponding IC50 value was determined and used to establish the experimental concentrations.

### Live staining and mitochondrial staining

PCLS were generated from 8–10-week-old NOD.Cg-*Prkdc^scid^*/J mice. Mitochondrial activity and cell viability was assessed using MitoSpy™ NIR DiIC1(5) (BioLegend, 424807) and Calcein AM (Life Technologies, L3224), respectively at 0, 1,3 and 7 days post PCLS generation. Briefly, PCLS were incubated with 50 nM MitoSpy dye and 2 µM Calcein AM in HBSS (Gibco, 14175-079) for 40 minutes at 37°C, then washed with HBSS and imaged. Positive control for death was obtained by treating PCLS with 70% methanol (Fisher, A452SK-4) for 1 hour at room temperature.

### Growth factor array

OS-17, OS186 and OS052 cells were seeded at 70-90% confluency then left in complete growth medium overnight. Cells were then washed twice using phenol-free DMEM/F-12 and supplemented with imaging media. After leaving the cells to incubate for 3 hours, the cells were either treated with AZD5991 (500 nM) or vehicle (media) and left to incubate for 4 hours. Supernatant was then collected, centrifuged at 1000 xG, and used at a 1:3 dilution for the Human Growth Factor Array L1 (Raybiotech, AAH-BLG-GRO-4) according to the manufacturer’s protocol. Glass slides were imaged using a Nikon Ti2 automated microscope.

### Sample preparation for proteomic analysis

OS-17 cells were seeded at 70-90% confluency then left in complete growth medium overnight. Cells were then washed twice using phenol-free DMEM/F-12 and supplemented with imaging media. Cells were either treated with vehicle (media), 500 nM MCL1 inhibitor (AZD5991), 100 nM MEK inhibitor (PD0325901) or 100 ng/mL FGF2 and left to incubate for 24 hours. After treatment, cells were detached with TrypLE, spun down and washed, once in phenol-free DMEM/F-12 and twice in PBS. Cells were then aliquoted and snap frozen for further processing.

Cell pellets were lysed and proteins denatured in 250 µL lysis buffer (4% sodium dodecyl sulfate (SDS), 50 mM Tris HCl, pH 8.0). Following incubation at 95°C for 10 min, samples were sonicated using a Q125 probe tip sonicator (QSonica, Newtown, CT) at amplitude 30% for three rounds of 30 s each. Lysates were cleared via centrifugation for 5 min at 20,000 RCF and supernatants were transferred to clean Protein LoBind tubes (Eppendorf, Framingham, MA). Proteins in each sample were quantified using the Pierce BCA Protein Assay Kit (Thermo Fisher Scientific, Waltham, MA). Aliquots of 100 µg protein per sample were precipitated in 3x volumes of ice-cold acetone overnight at 20°C. Precipitated proteins were recovered by centrifugation at 4°C, washed once in ice-cold acetone, and air dried at 4°C for 1.5 hrs.

Resulting protein pellets for each sample were solubilized in PreOmics Lyse buffer and processed to peptides using a PreOmics iST kit (PreOmics, Martinsfried, Germany) according to the manufacturer recommendations. Resulting peptides were dried down completely under vacuum at room temperature then reconstituted in LC-MS-grade water containing 0.1% formic acid (FA) (Fisher Scientific, Hampton, NH). Peptide yields were assessed using the Pierce Quantitative Fluorometric Peptide Assay (Thermo Fisher Scientific, Waltham, MA). Samples were diluted in 0.1% FA in water to accommodate injections of 400 ng peptide per sample for analysis by mass spectrometry (MS).

### Liquid chromatography-mass spectrometry (LC-MS) analysis

Peptides were analyzed by LC-MS on a Bruker timsTOF HT (Bruker Daltonics, Billerica, MA) MS platform coupled to an UltiMate 3000 HPLC with ProFlow (Thermo Fisher Scientific, Waltham, MA) facilitating inline nanoflow chromatography. Inline reversed-phase peptide separation was performed using a PepMap 5 mm Neo Trap Cartridge (Thermo Fisher Scientific, Waltham, MA), a PepSep C18 column (25 cm x 150 µm ID x 1.5 µm) (Bruker Daltonics, Billerica, MA), and a binary solvent system (A, LC-MS grade water, 0.1% FA and B, LC-MS grade acetonitrile, 0.1% FA) (Honeywell Chemicals, Charlotte, NC) delivered at a constant flow rate of 0.5 µL/min. Peptide separation was achieved using a linear gradient of 4-35% solvent B:solvent A over 60 min with a total MS acquisition time of 90 min. The MS front end leveraged a CaptiveSpray nanoflow electrospray ion source (Bruker Daltonics, Billerica, MA) containing a 20 µm Classic emitter (Bruker Daltonics, Billerica, MA) and equipped with column heating at 50°C (Analytical Sales and Services, Inc., Flanders, NJ). MS data collection was performed with dia-PASEF (data independent acquisition parallel accumulation-serial fragmentation) mode acquisition (1) using 22 dia-PASEF scans and 33 ion mobility windows with 25 Th width to cover a 0.6-1.45 V·s/cm2 (1/K0) ion mobility range and mass range of 397.3-1190.3 m/z. Capillary voltage was 1600 V. Ramp and accumulation times were set to 100 ms for a total cycle time of 2.44 s.

### Cell lysate derivation

OS-17, OS052 and OS186 wild type cells were plated at 70-90% confluency and were left to incubate overnight. Lysate was obtained the next day by scraping cells off the flask on ice, followed by repeatedly aspirating and pushing cell suspension through a 28-gauge needle. Suspension was then centrifuged at 14000 RPM at 4°C and the supernatant was snap frozen to be used for protein quantification and live cell imaging.

### Cell viability assay with dual Inhibitors

OS-17, OS052, OS186, and 143B wild-type cells were plated in a 96-well glass-bottom dish at a density of 7,000 cells per well, as described previously. The cells were then switched to imaging media and treated with AZD5991, as well as FGFR-targeting inhibitors—AZD4547, Dovitinib, Nintedanib, Erdafitinib, and Pazopanib—for 24 hours. After treatment, the cells were washed, fixed, and imaged. Cell counts were obtained using NIS software through nuclear segmentation.

### Fluorescent immunocytochemistry

OS-17, OS052 and OS186 wild type cells were plated in glass bottom 96 well plates and incubated as described for live-cell preparations, then treated with a titration of AZD5991 or FGF2 for 24 hours for Fra-1 measurement. Samples were fixed using 4% PFA, permeabilized with 0.1% Triton-X 100 and blocked with 3% BSA in PBS supplemented with 0.1% Tween-20. Primary antibody against Fra-1 (Santa Cruz Biotechnology, sc-28310, 1:400) and anti-mouse secondary antibody (Invitrogen, A21202, 1:400) were used to probe for Fra-1. Primary antibody against Nrf2 (Invitrogen, MA5-42371, 1:400) and anti-mouse secondary antibody (Jackson ImmunoResearch, 111-605-144, 1:400) were used to probe for Nrf2. Primary conjugated antibody (Santa Cruz, sc-136960, 1:400) was used to probe for HMOX1.

### Drug efficacy testing in experimental metastasis

To evaluate the therapeutic efficacy of dual MCL-1 and FGFR inhibition against established pulmonary metastases, NOD.Cg-*Prkdc^scid^*/J mice were inoculated via tail-vein injection with 1×10^6^ OS17 FLUC cells and monitored weekly by bioluminescence imaging (Xenogen IVIS Spectrum). At day 28 post-inoculation, mice were randomized into four equal groups to receive vehicle (10% DMSO, 40% PEG-300, and 5% Tween-80 in saline, 200uL IP, 3x/week), AZD5991 alone (60 mg/kg in 10% DMSO, 40% PEG-300, and 5% Tween-80 in saline, IP, 3x/week), pazopanib alone (30 mg/kg in 10% DMSO and 90% corn oil, PO BID, daily), or the combination for 2 weeks. Body weights and conditions were monitored twice weekly. Upon the first mouse reaching humane endpoints and showing confirmed pulmonary lesions, all animals were humanely euthanized. Lungs were blanched via right-ventricular saline perfusion and harvested for formalin fixation (largest lobe), OCT embedding (second largest lobe), and snap-freezing (remaining lobes).

### Drug efficacy testing in experimental metastasis for comparison with LungSITE

Thirty-four NOD.Cg-*Prkdc^scid^*/J mice were injected via tail-vein with single cell suspensions of 1 × 10^6^ OS17-FLUC cells. After 21 days, mice were anesthetized, imaged for luciferase activity via IVIS and randomized. Mice were divided among treatment groups: 7 receiving vehicle, 9 AZD5991, 9 Pazopanib and 9 AZD5991 and Pazopanib combination. Treatments were started at day 21 post injection. AZD5991 was dissolved in 40% polyethylene glycol (PEG)-300 (Fluka Analytica, 25322-68-3), 10% DMSO (Sigma-Aldrich, D2650), and 5% Tween-80 (Acros Organics, 9005-65-6) in sterile saline (VWR, GROW-MSDW-1000) as previously described (6). Pazopanib was dissolved in aqueous 0.5% hydroxypropylmethylcellulose (Sigma-Aldrich, H7509-25G) and 0.1% Tween 80, as described (56). Mice were treated with 60 mg/kg AZD5991 intraperitoneally on a Monday-Wednesday-Friday schedule for three doses. Mice were treated with 30 mg/kg Pazopanib twice daily via oral gavage for five doses. At day 26, mice were again imaged for luciferase activity via IVIS and euthanized at endpoint. Lungs were harvested and prepared for FFPE (Formalin fixed, paraffin embedded) sectioning. Lungs were sectioned into 5 µm sections using a microtome (Leica RM2125 RTS) and stained for vimentin to quantify metastatic burden.

### Fluorescent Immunohistochemistry

Slides were baked at 65°C for 30 minutes, then deparaffinized. Antigen retrieval was performed using citrate buffers (Sigma-Aldrich, C9999) and AR buffer (Agilent Technologies, CS70230-2) at in a pressure cooker set to 125°C and 90°C, respectively. Tissue slices were then blocked using 3% BSA for 30 minutes then stained with Vimentin antibody (Santa Cruz, sc-6260 AF790, 1:100). Slides were mounted using Diamond antifade mount then imaged using a Nikon Ti2 automated microscope

### Drug efficacy testing in LungSITE model for comparison with in vivo model

Metastasis bearing PCLS were transferred to a #1.5 glass bottomed 24 well plate and set up similarly to live-tissue microscopy plates. PCLS were treated with vehicle (imaging media), low dose AZD5991 (25 nM), pazopanib (2 µM) or combination of AZD5991 and pazopanib for 7 days and imaging media with included treatments were replenished every other day. PCLS were then fixed with 4% PFA and imaged.

### MATLAB analysis

Live-cell 2D imaging data and fluorescent immunohistochemistry images were analyzed using MATLAB R2025a (Mathworks, Natick, MA, USA). Custom MATLAB processing code (provided by the Michael, Pargett, Albeck Lab, UC Davis) was applied to imaging data to segment and track individual cells via the histone H2B nuclear fluorescence or Hoechst stain, measure ERKTR nuclear and cytoplasmic fluorescence intensities as well as antibody fluorescence intensities over time. At least two replicate experiments with 6000 cells per condition were used for each analysis.

### Mass spectrometry data processing and analysis

The DIA MS data files (Bruker .d) were searched against the UniProt human reference proteome (version from March 27, 2024, with isoforms included) using DIA-NN 1.8.1 (2) in library-free mode. Mass accuracy was set at 15 ppm and 20 ppm for MS1. Match between runs (MBR) was enabled to create a spectral library from the DIA runs and then used to reanalyze the data. Precursor FDR was set to 1%. Default settings were employed for all other parameters.

Data output was further analyzed in the R environment (version 4.5.1) (3) and with the following packages: tidyr (version 1.3.1), dplyr (version 1.1.4), ggplot2 (version 4.0.0), stringr (version 1.5.2), MSstats (version 4.16.1), and ggrepel (version 0.9.6). MSstats (4) was used for peptide-protein data processing and differential expression analysis with default settings employed for the ‘dataProcess’ function which encompasses normalization, summarization, missing value censoring, and imputation. Gene set enrichment analysis was performed using the GSEApy package in Python. Genes from differential expression results were ranked based on a composite score calculated as the product of fold change and the negative log10-transformed adjusted p-value. Pre-ranked GSEA was conducted using 1000 permutations against multiple gene set databases, including Gene Ontology Biological Process (GO-BP), Gene Ontology Molecular Function (GO-MF), KEGG, Reactome, and WikiPathways. Enrichment results were evaluated using normalized enrichment scores (NES) and statistical significance was determined using false discovery rate (FDR) q-values, where pathways with FDR q-values ≤ 0.05 were considered as significantly enriched.

### Drug synergy calculations

Synergyfinder.org was used to calculate Loewe synergy scores for dual treatment with AZD5991 and AZD4547. Software was also used to calculate standard deviation/error of mean.

### Spatial transcriptomics analysis

Four preprocessed 10X Visium datasets were downloaded from existing spatial transcriptomics data obtained from pretreated osteosarcoma lung metastasis (7) and data were analyzed in R (v 4.4.3) using the Seurat tools (Seurat v5.5.1 and SeuratObject 5.4.0). No additional quality control was realized. Cell type annotations were incorporated from the Budhathoki et al.(47) analysis and genes signatures calculated using the AddModuleScore function from Seurat, then spatially plotted with SpatialPlot. To improve visualization, image.alpha and min.cutoff were set to 0. The spot sizes vary between samples (OS1 to 1500, OS2 to 1400, OS4 to 1500 and OS6 to 2500). For analysis of FGFR1-HMOX1 overlap, the normalized data was extracted from the object. The spots were classified based on their values relative to the median score for FGFR1 and HMOX1 respectively.

## COMPUTATIONAL METHODS FOR SITE ANALYSIS

Live-cell osteosarcoma lung metastasis imaging data were analyzed using the Serial Imaging of Tumor and microEnvironment (SITE) computational pipeline ((30), https://github.com/cancer-dynamics/SITE_Murthy_2026). The analyses here used the same core workflow, with additions specific to the osteosarcoma lungSITE experiments and MCL1 inhibitor response studies outlined below. Tutorials and application of the data analysis and modeling methods applied here, including downloadable example data files, can be found at https://github.com/cancer-dynamics/SITE_Makkawi_2026.

### Cell segmentation and tracking

Osteosarcoma tumor cell cytoplasmic masks were generated using fine-tuned Cellpose models (57,58) trained on over 1300 manually annotated 2D images, including axial cross-sections, from live-cell imaging data. The trained model was then applied to full 3D time-lapse volumes to generate single-cell cytoplasmic masks for each frame. Nuclear masks were generated using a separately trained Cellpose model, trained on over 900 manually annotated 2D images, optimized for nuclear fluorescent signal and reporter-channel information. Reporter-cell identities were linked across time using the SITE optimal transport-based tracking framework. For tumors imaged in separate runs on multiple days, a global registration was performed before tracking single cells. Segmentation and tracking results were manually validated for accuracy across a random subset of 285 single-cell images and linkages between timepoints. Segmentation achieved an estimated 74.8% true-positive rate for complete tumor cell segmentation, defined as a segmentation mask correctly encompassing a single complete tumor cell, and a 98.9% true-positive rate for tumor cell identification, defined as a segmentation mask containing part of or a complete tumor cell. Over-segmentation was the most common error, with a segmentation mask containing an incomplete portion of a tumor cell in 21.6% of cases, while under-segmentation occurred in 3.5% of cases., defined as a segmentation mask containing more than one cell. ERK-KTR quantifiable masks were estimated to be present in 78.3% of cases, defined as segmentation mask containing sufficient nuclear and cytoplasmic portions to confidently quantify the C/N ratios. 91.5% of tumor cell masks were appropriately tracked between consecutive frames. The resulting tracked cytoplasmic masks provided the basis for extracting longitudinal single-cell trajectories of ERK activity, FIRE activity, position, motility, contact state, and survival or death fate.

### Reporter activity quantification

ERK activity was quantified at the single-cell level from ERK kinase translocation reporter fluorescence. For each segmented cell, mean reporter intensity was measured in nuclear and cytoplasmic compartments, with the cytoplasmic compartment defined by subtracting the nuclear mask from the full cytoplasmic cell mask. Background fluorescence was estimated from non-cellular regions in each frame and subtracted before ratio calculation. ERK-KTR activity was then quantified as the cytoplasmic-to-nuclear fluorescence intensity ratio, with increased cytoplasmic localization corresponding to increased ERK activity. FIRE expression was calculated as the average intensity inside the nuclear compartment of each cell.

### Cluster size description

To characterize the size of reporter cell clusters, connected regions in the ERK-KTR intensity volumes were extracted. At each timepoint, the ERK-KTR intensity channel was first smoothed using a 3D Gaussian filter (σ = 10 µm in the lateral dimension, adjusted by z-scale axially), and a z-score– normalized foreground mask was generated using a threshold of z = 2.0. Connected components within this foreground mask were labelled to define multicellular clusters. Labelled cytoplasmic segmentations were then assigned to ERK-KTR-defined clusters by intersecting reporter cell labels with each connected region.

### Cell local environment proximity

To quantify local tumor–microenvironment organization, we calculated the spatial relationships around a single-cell boundary to its nearest cell or tissue boundary. 3D boundary maps were computed for each segmented reporter cell at each time point, and lung tissue and non-tumor microenvironmental compartments were segmented using ilastik-based semi-supervised random forest classification. These masks provided the spatial context required to classify tumor-cell boundary proximity with neighboring tumor cells, host lung tissue, and extracellular or non-cellular regions. Cell and tissue surfaces were sampled at 1.0 µm boundary resolution, and each boundary point was assigned nearest-neighbor relationships to tumor-cell, host-tissue, and extracellular compartments. We defined the proximity p to a cell/surface as the inverse mean distance around the boundary to neighboring compartments, 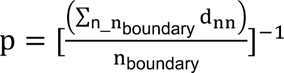 with n^n^_boundary_ the number of boundary points of the cell, and dnn the nearest neighbor distance from each boundary point to it’s nearest neighboring cancer cell, or host cell boundary point. The inverse mean distance is a natural scaling of spatial context where a cell was completely surrounded on the boundary by neighboring cancer cells has a proximity to cancer cells of p = 1.0 µm^−1^ (with 1.0 µm boundary resolution), while an isolated cell will tend to p = 0.0 µm^−1^. We use these proximities as a local measure of the cellular environment, and to visualize the environmental dependence of drug response.

### Local environment proximity featurization

To quantify the spatial organization of the local tumor microenvironment around each cell, we represented neighboring cancer-cell and host-cell surfaces as boundary-resolved proximity fields and summarized these fields using a multipole expansion. This approach was motivated by the approximation that the steady-state concentration profile arising from a localized sender decays as 1/r, so that inverse distance can be interpreted as a local potential-like measure of environmental influence. Consistent with this framework, the SITE methodology captures the spatial and temporal influence of cell-cell interaction using this proximity-based characterization of local environmental influence.

Boundary maps were first computed for each segmented cell at 1.0 µm resolution, yielding sampled boundary points across the 3D cell surface and nearest-neighbor surface assignments to surrounding compartments. This follows the same boundary-based representation used in the SITE platform to quantify tumor–microenvironment interactions from segmented cell surfaces (30). For each boundary point k on cell i, and for each neighboring compartment type c (surrounding cancer cells or host tissue), we computed the nearest-neighbor distance 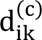 from that boundary point to the corresponding compartment surface. Local proximity was defined as the inverse distance,

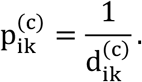

To center and rescale this proximity signal before multipole decomposition, we assigned each boundary point a scalar boundary “charge”

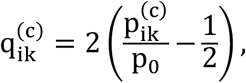

where p_-_ = 1/Δb and Δb is the boundary sampling resolution (here, 1.0 µm).

With this definition, a neighboring surface in contact lying approximately one boundary-resolution unit away gives q ≈ 1, whereas a surface which is a long distance away tends to q ≈ −1. In practice, this transformed inverse-distance field serves as a boundary-distributed approximation to the local potential generated by the surrounding compartment.

The spatial distribution of 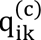 over the cell surface was then decomposed using a spherical multipole expansion evaluated on the discrete set of boundary points. For multipole order *_l_*, the complex moments M*_P_*_/_were computed over all boundary points and summarized as a normalized magnitude,

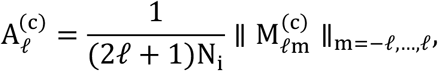

where N_i_is the number of sampled boundary points for cell i. We retained the *_l_* = 0 and *_l_* = 1 terms for downstream analysis. The *_l_* = 0 term captures the isotropic component of local environmental proximity, corresponding to the overall mean strength of nearby surrounding cancer-cell or host-cell surfaces. The *_l_* = 1 term captures the first anisotropic component, reflecting whether that environmental proximity is polarized to one side of the cell.

These multipole features were computed separately for the distributions of surrounding cancer cells and surrounding host cells, yielding low-dimensional descriptors of each cell’s local microenvironment. In this way, each cell was featurized not only by how close it was, on average, to neighboring compartments, but also by whether those neighboring compartments were spatially symmetric or directionally biased around the cell boundary.

### Death identification

We found that many death events could be validated via monitoring changes in cell morphology, as cell deaths under MCL1i led to characteristic rupture and identifiable apoptosis, see **Supplementary Figure 2B**. Linked cell trajectories were pruned with a cutoff optimal transport cost earth movers distance (cite https://jmlr.org/papers/v22/20-451.html) of 10.0 µm, and single-cell trajectory endpoints manually inspected by evaluating maximum axial projection images of the ERKTR cell over a 2 hour window from the trajectory endpoint. Indices of all identified cell deaths in our single-cell trajectories were stored for further analysis linking ERK signaling to drug response and cell fate. Death rates over time were estimated from the slope of the cumulative distribution of death events over time in each condition.

### Spatiotemporal correlation between cell death and ERK signaling

To quantify how cell death events were associated with subsequent ERK activation in nearby cells, we computed a death-to-signal spatiotemporal correlation function from tracked single-cell trajectories. ERK activity was binarized into ERK ON (C/N ratio>1.0) and ERK OFF states (C/N ratio<1.0). For each death event, the dying cell was treated as the source event at time t, and ERK signaling states of neighboring cells were measured at a later time t + Δt. Distances were computed from the position of the dying cell at the source frame to the positions of all tracked cells at the lagged frame, after rescaling the axial dimension to account for anisotropic voxel spacing. Pairwise distances were binned into radial intervals, and for each radial bin the mean ERK activity of neighboring cells was calculated at each time lag. Operationally, this yielded a radial, time-lagged death-to-signal correlation function of the form

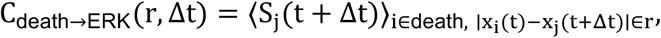

where S_j_(t + Δt) denotes the probability of finding an ERK ON neighboring cell j at lagged time t + Δt, and the average is taken over all cell pairs in which cell i was annotated as a death event at time t and the pairwise separation fell within radial bin r.

In practice, death events were identified from the tracked single-cell trajectories, and the correlation analysis was performed over all imaging frames within the selected analysis window. For each frame containing one or more death events, all dying cells in that frame were used as source cells. The ERK state of neighboring cells was then evaluated after a specified lag Δt, and averaged within radial bins. The resulting 2D map reports the probability or mean fraction of ERK-active cells as a function of distance from a death event and elapsed time after that event.

To generate replicate-level summaries, the death-to-signal correlation was computed separately for each experiment or imaging model and then averaged across replicates. Radial bins extended from short-range nearest-neighbor distances out to the maximum interaction radius used in the analysis, and time lags were evaluated over successive imaging intervals. Axial distances were rescaled before binning so that radial distances reflected physical spacing in microns rather than voxel units.

As a null comparison, we randomly shuffled cells identified as death events, with other cells within the same frame. Comparison to null models (**Supplementary Figure 2E**) allowed us to assess whether local ERK activation near death events exceeded that expected from the global fraction of ERK-active cells alone and validate a time- and distance-dependent increase in ERK activity surrounding death events, consistent with a local adaptive signaling response to nearby cell death.

### Death event density metric and ERK activity transition probability

To quantify how nearby cell death events influenced the probability of ERK state switching in surviving cells, we defined a local death-exposure metric analogous to the local signaling-density framework used for ERK–ERK interactions (cite Murthy2026). The rationale was that, if cell death releases locally acting bioactive factors, then the influence of multiple death events on a given cell should scale with both the number of nearby death events and their distance from that cell, with closer events contributing more strongly. As in related SITE analyses, this influence was modeled using an inverse-distance weighting, consistent with a steady-state sender field that decays approximately as 1/r.

For each tracked cell i at time t, we computed a death-proximity score S_d_/R by summing the inverse-distance-weighted contributions of nearby death events occurring within a defined retrospective time window. This metric was defined as

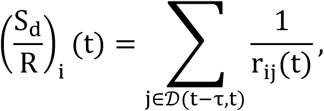

where _D_(t − τ, t) denotes the set of cells that underwent death within the preceding time window of duration τ, and r_ij_(t)is the distance between cell i and death event j. In the primary analysis, τ was set to 8 hours, such that S_d_/R captured the cumulative recent local exposure of a cell to nearby death events.

Cells were classified into ERK-KTR_low_ and ERK-KTR_high_ states. For each tracked cell, ERK state transitions between consecutive frames were then enumerated, yielding OFF→ON and ON→OFF transition events. Transition probabilities were stratified as a function of the death-proximity score S_d_/R, allowing us to quantify whether recent local exposure to cell death altered the probability of ERK activation or deactivation.

Operationally, cells were grouped into bins of low to high S_d_/R, and within each bin, the probability of ERK activation was calculated as

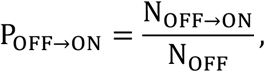

where N_OFF→ON_is the number of cells that transitioned from ERK-KTR_low_ to ERK-KTR_high_ between consecutive frames, and N_OFF_ is the total number of ERK-KTR_low_ cells eligible for such a transition in that bin. Similarly, the ERK deactivation probability was calculated as

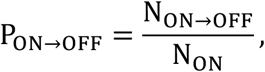

where N_ON→OFF_ is the number of ERK-KTR_high_ to ERK-KTR_low_ transitions and N_ON_ is the total number of ERK-KTR_high_ cells in the corresponding bin.

This analysis yielded a data-driven description of how local death exposure biased ERK state dynamics. Increased S_d_/R indicated greater recent proximity to death events and therefore stronger putative exposure to death-associated local signals. By comparing ERK transition probabilities across S_d_/R bins, we assessed whether nearby death events preferentially promoted ERK activation.

### Contact niche states to map microenvironmental dependence

To summarize local tumor microenvironment organization at the single-cell level, we defined contact niche states from features describing each cell’s local proximity to surrounding cancer cells and host tissue. These features were derived from boundary-resolved measurements of the local environment (see methods: Local environment proximity featurization) and then grouped using our previously described trajectory-embedding and kinetic-clustering framework developed for data-driven cell-state discovery (cite https://www.sciencedirect.com/science/article/pii/S2405471226000499).

Briefly, for each tracked cell and time point, we computed the l=0 and l=1 moments of the local cancer-cell and host-cell proximity. Single-cell trajectories through this feature space were then used to construct a data-driven dynamical embedding (for single-timepoints, or trajectory length=1), and kinetically similar microstates were coarse-grained into metastable contact niche states from local tumor–microenvironment interaction features over time.

For the present study, this procedure yielded three reproducible contact niche states that were readily interpretable based on the degree of surrounding cancer-cell proximity: low cancer proximity (LCP), mixed cancer proximity (MCP), and high cancer proximity (HCP). These states corresponded to progressively greater enrichment of neighboring cancer-cell influence within the local microenvironment and were used as a compact description of each cell’s spatial niche during MCL1 inhibitor response. This contact niche state representation was used downstream to stratify ERK signaling behavior during drug response. By reducing complex local spatial organization into a small number of metastable, dynamically informed categories, the contact niche framework enabled integration of microenvironmental context with signaling and fate measurements at the single-cell level.

### Composite state-space construction for adaptive drug-response modeling

To model how local microenvironment, death exposure, ERK signaling state, and cell fate jointly evolved during MCL1 inhibitor response, we constructed a discrete composite state space from single-cell trajectory features. This analysis extended the trajectory-informed contact niche framework used above to construct Markov models which described the signaling-dependent state transitions over time.

The base state of each cell was its contact niche assignment, coarse-grained into three metastable categories: high cancer proximity (HCP), mixed cancer proximity (MCP), and low cancer proximity (LCP). To this base niche state, we appended three additional binary variables: alive versus dead, low versus high death-proximity exposure, and ERK-KTR_low_ versus ERK-KTR_high_ signaling state. ERK state was defined from the single-cell ERK-KTR cytoplasmic-to-nuclear ratio using the same binarization threshold (c/n=1.0) used elsewhere in the signaling analyses. Death-proximity state was defined from the S_d_/R window 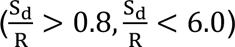 where increased ERK-OFF -> ERK-ON was observed in the ERK activation probability analysis (**Figure 3C**), capturing the subpopulation with recent exposure to nearby death events likely to trigger increased ERK signaling.

Each tracked trajectory step was therefore assigned to one of 3 × 2 × 2 × 2 = 24 coarse-grained composite states, corresponding to all combinations of contact niche state, life state, death-proximity state, and ERK signaling state. This yielded a compact discrete representation of the adaptive drug-response landscape in which cells could be followed through changes in microenvironmental niche, signaling status, recent local death exposure, and eventual survival or death.

### Transition matrix estimation

Transition probabilities between composite states were estimated directly from tracked single-cell trajectory steps. For each consecutive trajectory pair (x_t_, x_t+1_), the cell’s feature vector at time t and time t + 1 was mapped to the nearest composite-state center, and transition counts between all coarse-grained states were accumulated to form an empirical transition matrix, with the lag time of the model therefore matching the imaging step used for trajectory extraction.

In practice, the composite-state centers were built hierarchically by augmenting the contact niche state representation with additional discrete coordinates for alive/dead status, low/high S_d_/R, and low/high ERK state. Death status for each trajectory step was assigned by identifying whether the destination cell at t + 1belonged to the curated set of death events. S_d_/R values were taken from the death-proximity calculations described above, and ERK state was assigned from the most recent finite ERK-KTR value associated with that trajectory step.

### Death rate estimation

Because the number of observed death transitions was limited relative to the number of alive-to-alive transitions, death rates were estimated separately from the coarse-grained transition counts and then re-imposed on the transition matrix. Specifically, the empirical probability of death among ERK-KTR_high_ and ERK-KTR_low_ cells was calculated directly from trajectory steps as

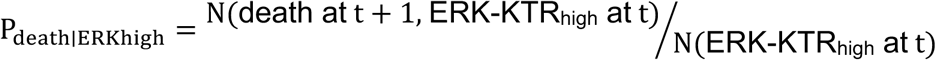

and

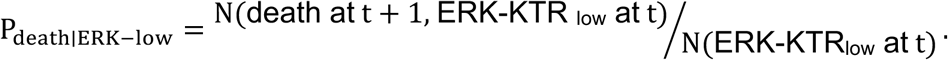

These ERK-stratified death rates were then inserted into the composite transition matrix by assigning each alive ERK-KTR_high_ state a transition probability into its corresponding dead ERK-KTR_high_ state, and likewise for ERK-KTR_low_ states. Dead states were treated as absorbing states. In the implemented model, transitions from high S_d_/R states to corresponding low S_d_/R states were suppressed and reassigned to the matched high-S_d_/R state, reflecting the intent to preserve recent death exposure as a cumulative local variable over the modeled time window rather than allowing it to decay immediately in a single Markov step.

Since our manually curated set of captured death events was not complete of every cell death in the imaged tumors, we fit an overall multiplicative value to both ERK-ON and ERK-OFF death rates to the overall tumor cell number as a function of time, with a good fit obtained by multiplying death rates by 3.15, suggesting our manually curated set of death events captured about 1/3 of death events overall in the tumor.

### ERK-switching rate estimation

From the fitted transition matrix, several interpretable quantities were extracted. First, ERK switching probabilities were computed separately for cells in low- and high-death-exposure environments. For each contact niche state, the ERK activation probability under low S_d_/R was defined as the summed probability of transitions from alive ERK-KTR_low_/Sd/R_low_ states into alive ERK-KTR_high_ states, with the corresponding ERK deactivation probability defined analogously from alive ERK-KTR_high_/ Sd/R_low_ states into alive ERK-KTR_low_ states. The same calculations were repeated for Sd/R_high_ states. This yielded niche-specific estimates of P_OFF→ON_ and P_ON→OFF_ under low and high death exposure. In parallel, ERK-dependent death probabilities were computed for each contact niche state by summing transition probabilities from alive ERK-KTR_low_ or alive ERK-KTR_high_ states into any dead state. This formulation directly quantified the dependence of death risk on signaling status within each contact niche.

### Validating transition model estimation

Observed state probabilities over time were calculated independently from the raw imaging data for comparison with model evolution. At each time point, all tracked cells were assigned to the composite state space, and the fraction of cells in each contact niche state was calculated either after pooling over ERK and S_d_/R sub-states or after stratifying by ERK-KTR_low_ and ERK-KTR_high_ populations. These observed state fractions were evaluated in sliding 1-hour windows and scaled by the relative surviving cell number, so that the time-dependent state occupancies reflected both redistribution among states and overall population loss during treatment.

Model-predicted state probabilities were generated by initializing the Markov model from the empirical state distribution at the start of the experiment and iteratively propagating that probability vector forward by repeated multiplication with the fitted transition matrix. For the main MCL1 inhibitor simulations, the initial state distribution was obtained from the earliest tracked time window after treatment onset. Predicted occupancies of HCP, MCP, and LCP states were then compared directly with the observed state fractions over time, both for the total alive population and for ERK-KTR_low_ and ERK-KTR_high_ subpopulations.

To estimate the effect of ERK inhibition in silico, we modified the fitted transition matrix to block ERK activation transitions. Operationally, all transitions from alive ERK-KTR_low_ states to alive ERK-KTR_high_ states were set to zero, and the affected rows were renormalized so that the remaining transition probabilities summed to one. The initial condition was also projected into the ERK-KTR_low_ manifold by transferring any probability mass in alive ERK-KTR_high_ states into the corresponding alive ERK-KTR_low_ states. This procedure produced an idealized “complete ERK inhibition” perturbation model in which ERK-KTR_high_ states could not be newly acquired. The resulting simulated state occupancies and total surviving population were then compared with the unperturbed MCL1 inhibitor model to estimate how blocking ERK-KTR_low_ to ERK-KTR_high_ transitions would alter adaptive resistance and overall cytotoxicity. Uncertainty in the estimated transition-derived quantities was assessed by bootstrap resampling at the imaging-set level. In the notebook implementation, replicate imaging sets were resampled with replacement, and transition probabilities, ERK switching rates, death rates, and observed state occupancies were recalculated across bootstrap replicates to obtain mean estimates and confidence intervals.

### Confidence Interval Estimation

Confidence intervals for Figures 2C, 3F, 3G, 5E, Supplementary figures 2F, 6B, 7C and 7D were estimated by bootstrapping across imaging fields of view (1000 bootstraps) using a Bayesian bootstrapping procedure (cite https://pubs.acs.org/doi/full/10.1021/acs.jctc.9b00015). Additionally, Figure 3C bootstrapped over biological replicate variation, with each bootstrapped sample taken leaving one replicate out (*_n_* = 100 per replicate).

## SUPPLEMENTARY FIGURES

**Supplementary Figure 1.**
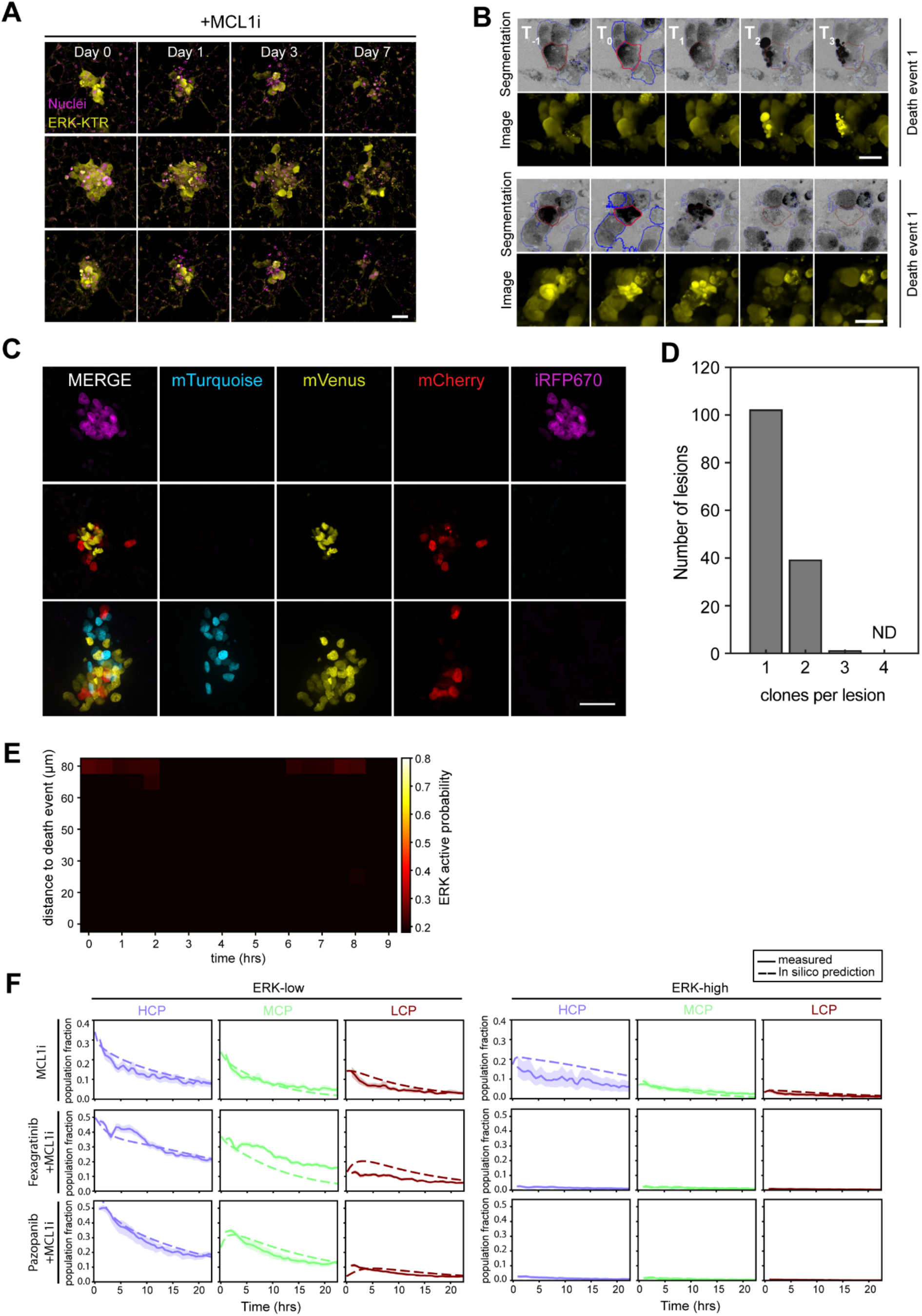
**(A)** 3D rendered images (top) of PCLS showing OS-17 ERKTR (yellow) integrated in the lung parenchyma (grey) and Hoechst-stained nuclei (blue), along with maximum intensity projection images of OS-17 ERKTR 0, 3 and 7 days post generation of PCLS. Magenta arrows indicate cell movement, white arrows indicate ERK state swutch and blue arrows indicate cell division. Scale bar = 50 µm. **(B)** Maximum intensity projection images of Calcein AM (green) and MitoSpy staining (magenta) of precision cut lung slices 0, 3 and 7 days post generation and culturing ex vivo. Methanol treated PCLS is included as a negative control. Scale bar =100 µm. **(C)** 3-D rendering of PCLS (top) showing ERK-KTR reporter cells (yellow) in the lung parenchyma (grey) along with 3-D rendering of segmentation (bottom) of the same reporter cells and lung parenchyma. Scale bar = 50 µm.

**Supplementary Figure 2.**
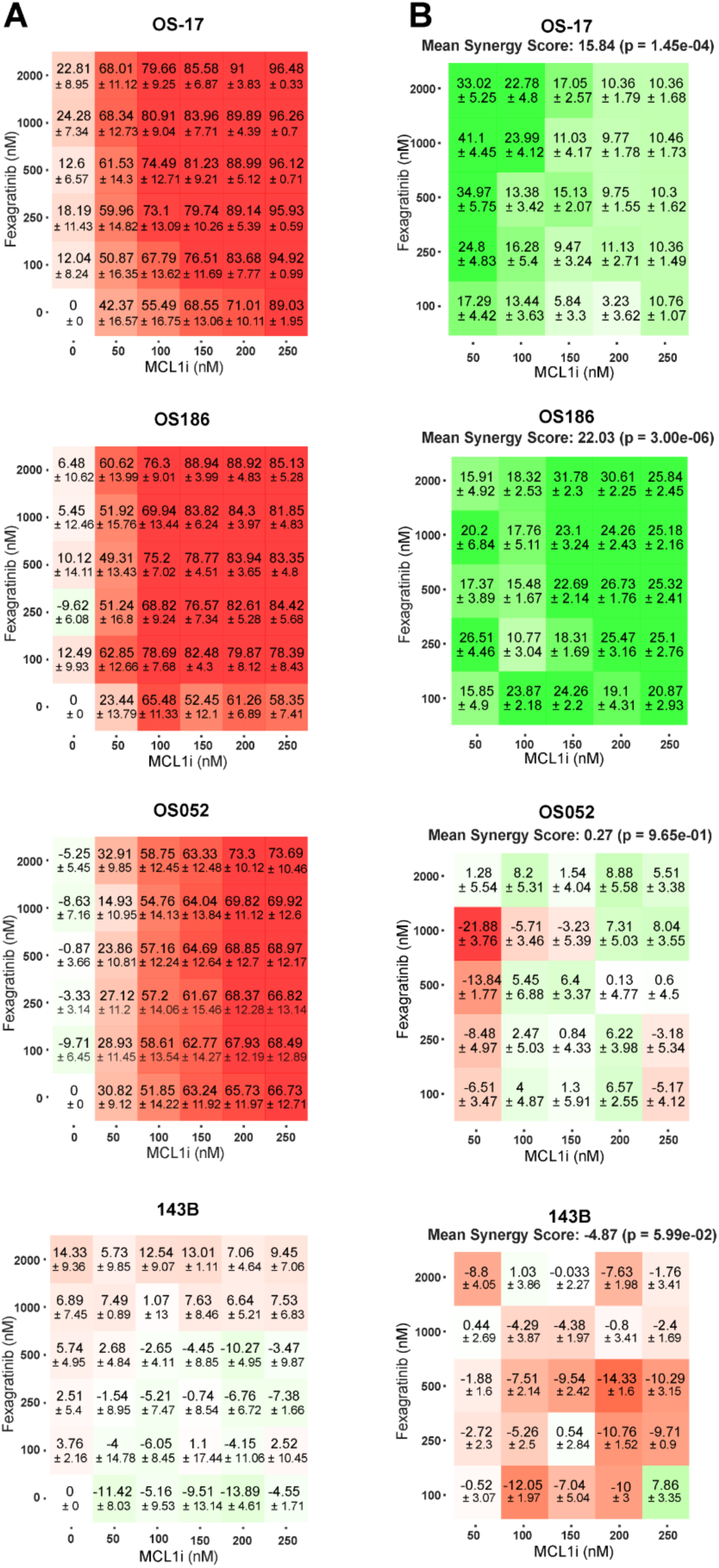
**(A)** Representative maximum intensity projection images of OS-17 reporter cells in LungSITE at the indicated time points after treatment with 500nM MCL1 inhibitor, AZD5991. Images show ERK-KTR (yellow) and nuclei, indicated by both H2B reporter and Hoechst stain (magenta). Scale bar = 50 µm. **(B)** Images showing cell death event identification using the SITE computational pipeline. Top panels show cell segmentations overlaid on maximum-intensity projection images generated by the pipeline. The bright red outline indicates the selected cell at the time frame immediately preceding cell death, while dark red outlines indicate the same cell across time frames before and after death identification. Blue outlines indicate the segmentations of neighboring cells. Corresponding maximum-intensity projection images obtained using NIS-Elements software are shown in the bottom panels. Scale bar = 25 µm. **(C)** Representative maximum intensity projection images of OS-17 H2B reporter cells in LungSITE showing mono- and poly-clonality. mTurquoise, mVenus, mCherry and iRFP670 H2B reporters were used. Scale bar = 50 µm. **(D)** Bar chart showing the distribution of the number of H2B clones per OS-17 lesion in LungSITE. Data was collected from 2 replicate experiments. **(E)** ERK activation probability heatmap as a function of distance to a cell death event (µm) and time of a null model, where cells identified as death events were shuffled with other cells within the same time frame. **(F)** Graphs showing experimentally measured (solid lines) and in silico predicted (dashed lines) changes in contact niche state population fractions, stratified by ERK_high_ and ERK_low_ states, over time after treatment of OS-17 cells LungSITE with 500nM MCL1i, or 100nM MCL1i in combination with 2µM fexagratinib and pazopanib. Shaded areas represent the bootstrapped 95% confidence interval (CI). Data was collected from 3 replicate experiments.

**Supplementary Figure 3.**
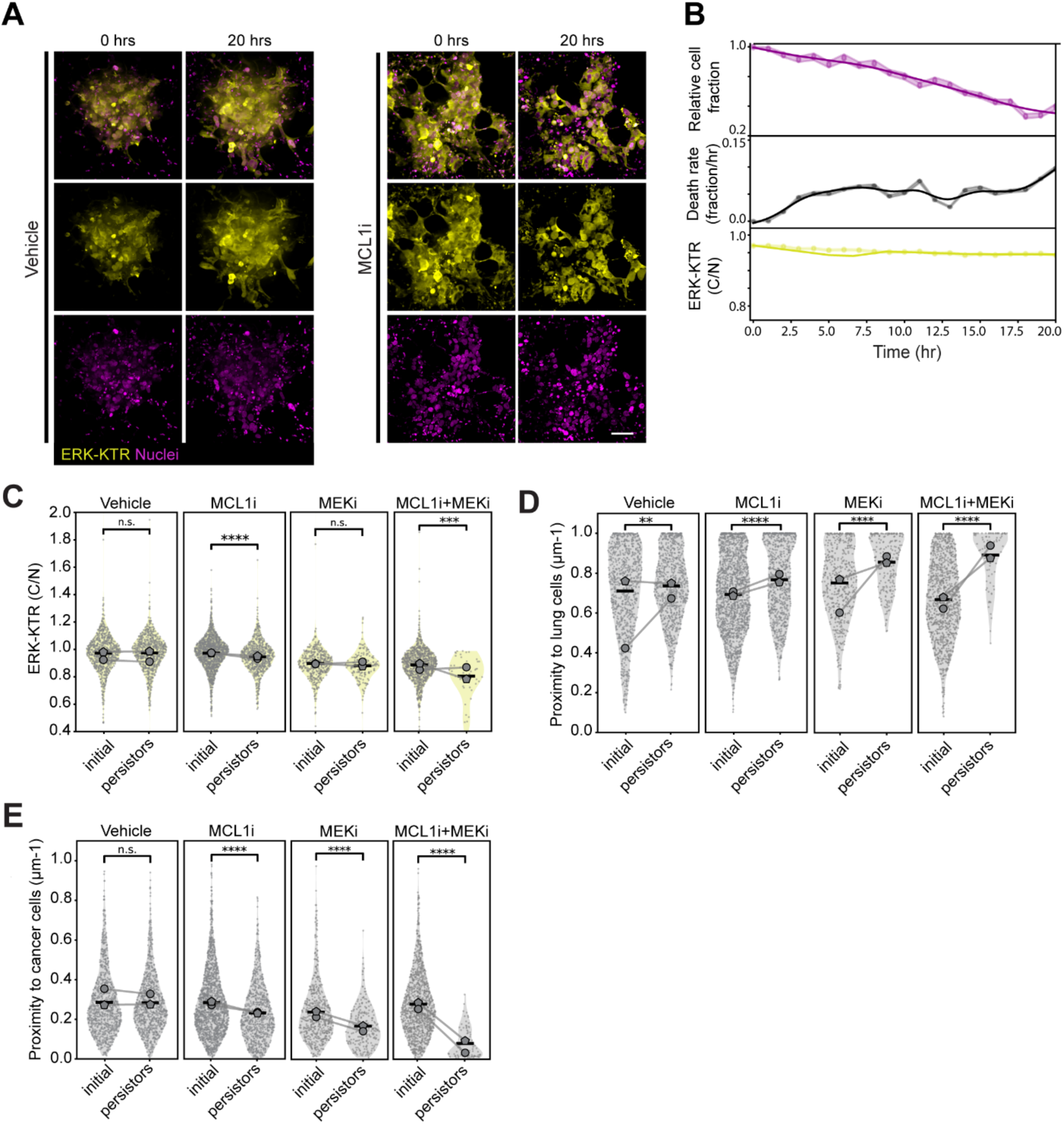
**(A)** Heatmaps showing percentage of cell death in OS-17, OS186, OS052 and 143B cells treated for 24 hours with titrations of fexagratinib (y-axis) and MCL1i, AZD5991 (x-axis). Average counts and standard deviation from 4 independent replicate experiments are shown. **(B)** Heatmaps showing Loewe synergy scores for data shown in (A). Overall score is included on top of heatmaps, and individual scores for specific concentration combinations and standard deviations are included as well.

**Supplementary Figure 4.**
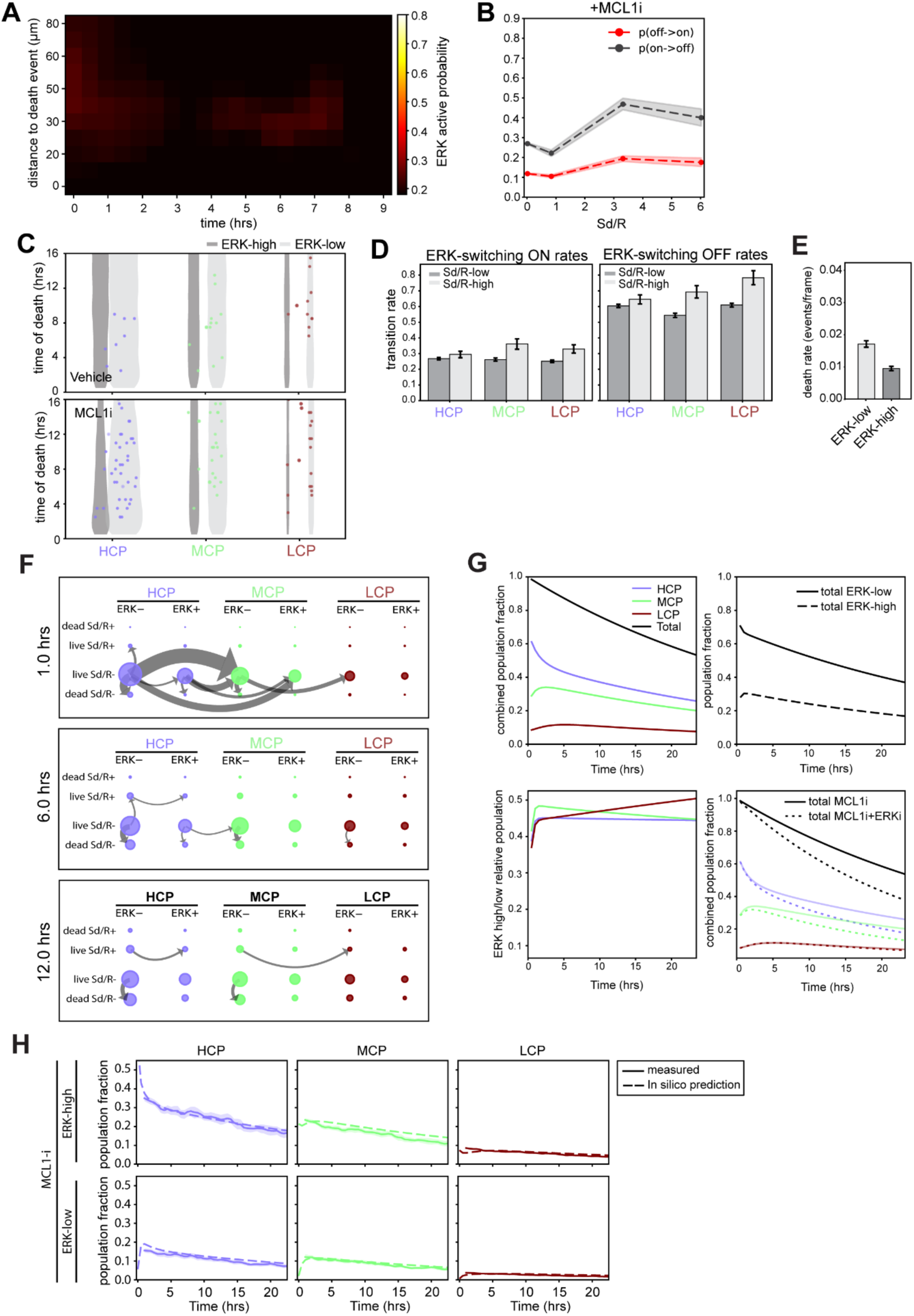
**(A)** Representative maximum intensity projection images of143B reporter cells in LungSITE at the indicated time points after treatment with Vehicle or 500nM MCL1 inhibitor, AZD5991. Images show ERK-KTR (yellow) and nuclei, indicated by both H2B reporter and Hoechst staining (magenta). Scale bar = 50 µm. **(B)** Line graphs depicting remaining cell fraction over time (top, magenta line), death rate as fraction per hour (middle, black line), and ERK-KTR activity as a function of C/N ratio (bottom, yellow line). Data was obtained over the indicated time course following treatment with 500nM MCL1 inhibitor, AZD5991, and is the aggregate of 2 biological replicates. Shaded areas represent the bootstrapped 95% confidence interval (CI). **(C)** Violin plots depicting ERK-KTR activity (C/N ratio) of 143B cells in LungSITE at treatment initiation and 20hrs post-treatment with vehicle, 500 nM MCL1 inhibitor, 100 nM MEK inhibitor or a combination of MCL1and MEK inhibitors. Mean of separate experimental replicates are indicated by grey symbols, the aggregate mean is indicated by the horizontal line. The connecting line between symbols represents the initial and persister activities of each replicate. **(D)** relative proximity of an individual living cancer tothe lung, or **(E)** relative proximity of an individual living cancer cell to other cancer cells at treatment initiation and 20 hours post-treatment. Relationships were extracted from the datasets presented in panel (B).

**Supplementary figure 5.**
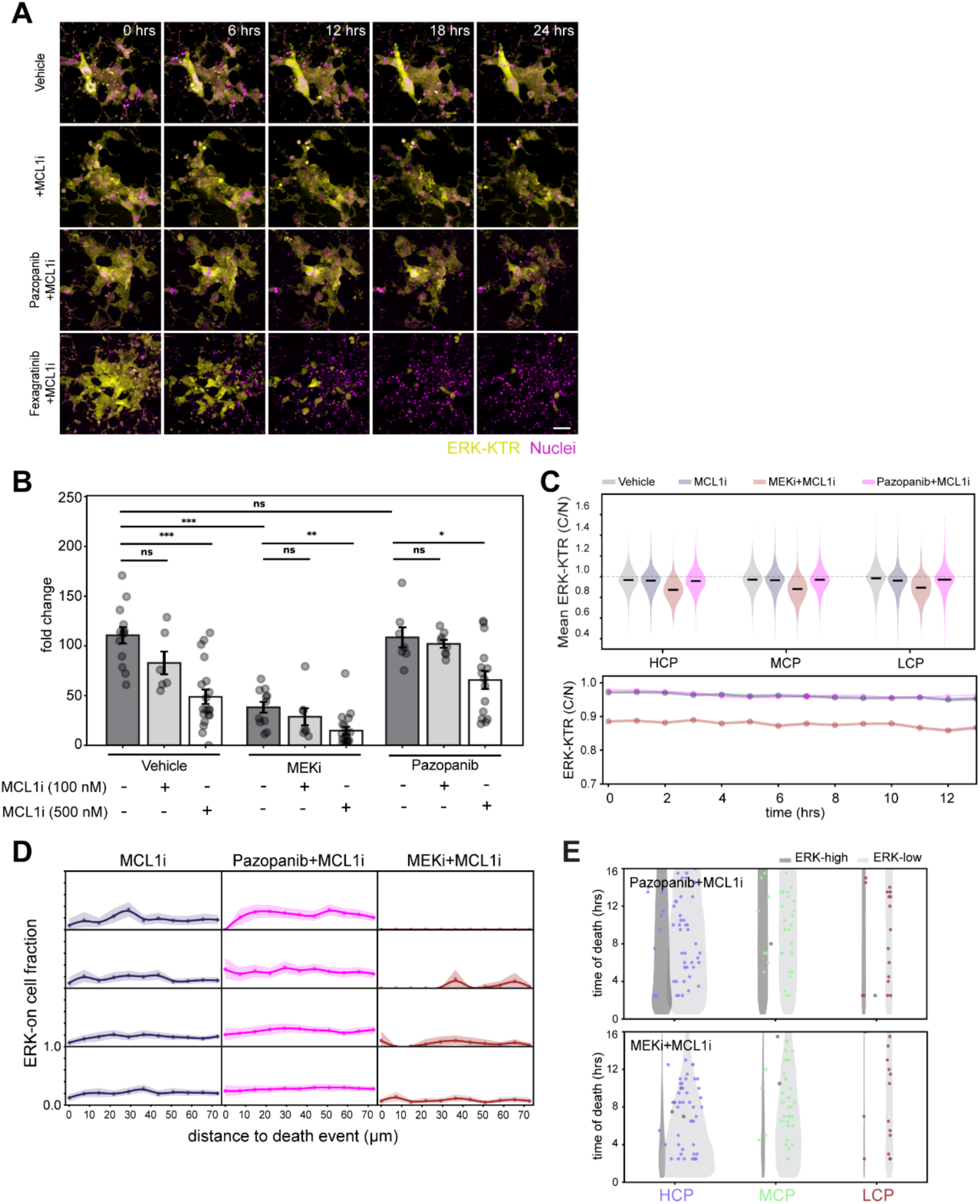
**(A)** ERK activation probability heatmap as a function of distance to a cell death event (µm) and time using data obtained from LungSITE metastatic 143B lesions treated with a 500nM MCL11i. Data is obtained from 2 biological replicates. **(B)** Calculated ERK state probably for ERK activation, off → on (red) and ERK deactivation, on ◊ off (black) as a function of proximity to death events (Sd/R) in 143B LungSITE metastasis treated with 500nM MCLi. **(C)** Violin plots showing time of death of ERK_high_ and ERK_low_ 143B cells in LungSITE under Vehicle and 500nM MCL1i conditions, stratified by niche states. **(D)** Bar plots showing rates of ERK switching in tumors treated with 500nM MCL1i, stratified by proximity to death events (Sd/R) and contact niche states; high, mixed, and low cancer proximities. **(E)** Bar plot showing average death rates of ERK_low_ and ERK_high_ tumor cells following treatment with 500nM Mcl1i in LungSITE. **(F)** Markov model data demonstrating flux between HCP to MCP and LCP contact niche states 1, 6, and 12 hrs following treatment in LungSITE. **(G)** Graphs showing experimentally measured changes in contact niche state population fractions over time (top-left), fraction of ERKhigh and ERKlow cells (top-right) over time, and relative ERKhigh/low populations (bottom-left) over time after MCL1i treatment in LungSITE. The graph bottom-right depicts in silico prediction of changes in fraction of contact niche states over time (dotted lines) under dual inhibition by ERKi and MCL1i superimposed on experimentally collected data (solid lines).**(H)** Graphs showing experimentally measured (solid lines) and in silico predicted (dashed lines) changes in contact niche state population fractions of 143B cells, separated into ERK_high_ and ERK_low_ cells, over time after treatment in Lung SITE following treatment with 500nM MCL1i. Shaded areas represent the bootstrapped 95% confidence interval (CI).

**Supplementary figure 6.**
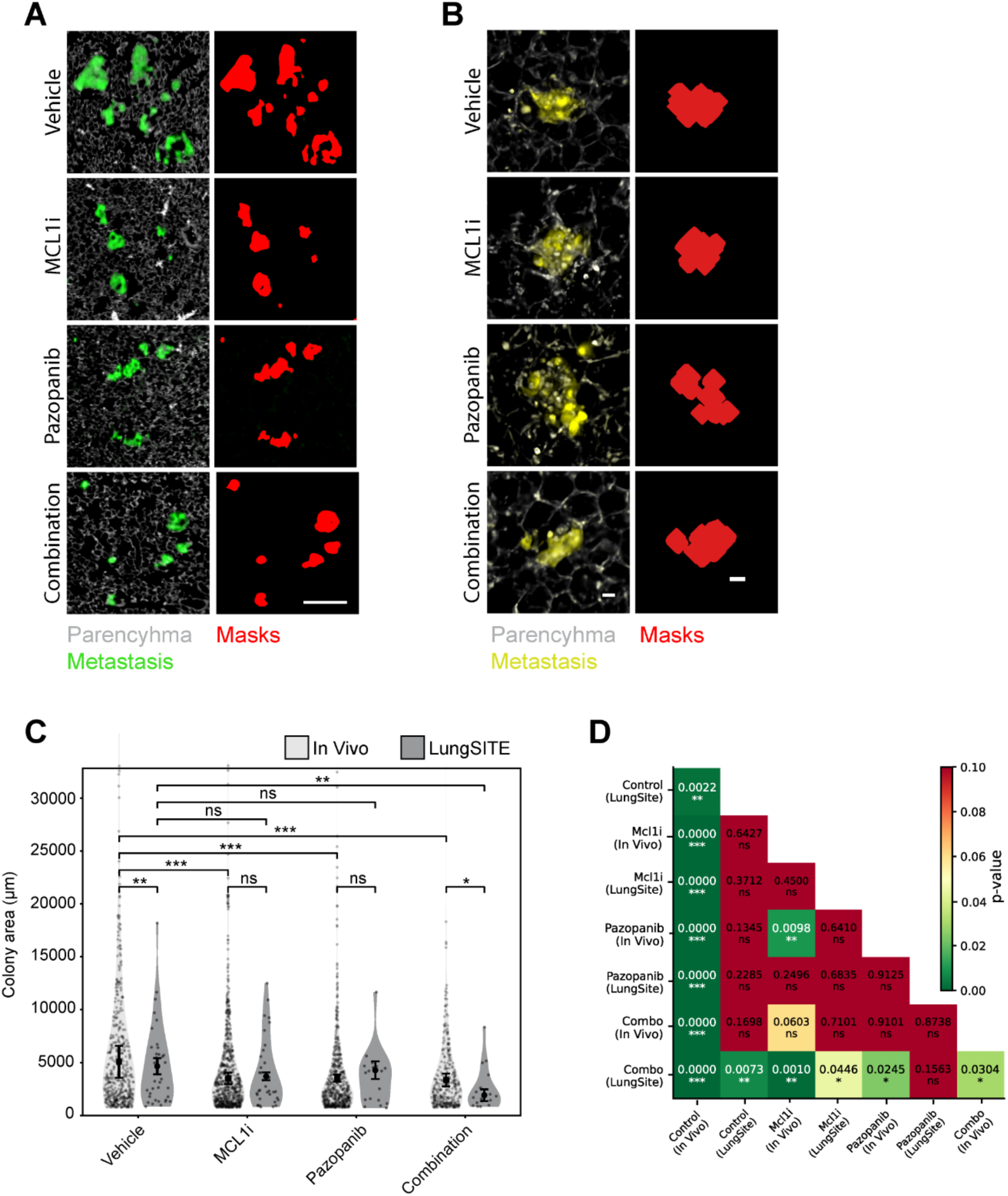
(**A)** Representative maximum intensity projection images of 143B cells, ERK-KTR (yellow) and nuclei (magenta) indicated by both H2B reporter and Hoechst stain, in LungSITE model treated with 500nM MCL1i alone and in combination with 2µM pazopanib or 100nM MEKi are shown. Scale bar = 50 µm. **(B)** Bar plot showing fold-change in cell count of 143B cells in LungSITE following 24hrs exposure to the indicated drug concentrations. Data was aggregated from 2 independent replicate experiments. **(C)** Violin Plot (Top) showing distribution of mean ERK-KTR activity (C/N ratio) of tumor cells in LungSITE following exposure to Vehicle, 500nM MCL1i, or 2µM pazopanib or 500nM MEKi in combination with 100nM MCL1i stratified by contact niche states. Line plot (bottom) comparing ERK-KTR activity with the indicated time-period following treatment with 500nM MCL1i, alone or in combination with 2µM pazopanib or 100nM MEKi. Shaded areas represent the bootstrapped 95% confidence interval (CI). **(D)** Line plots showing mean fraction of “ERK ON” cells at different time points as a function of distance from neighboring death events following treatment with 500nM MCL1i, alone or in combination with 2µM pazopanib or 100nM MEKi. Shaded areas represent the bootstrapped 95% confidence interval (CI). **(E)** Violin plots of time of death for ERK-high and ERK-low 143B cells in LungSITE in response to 2µM pazopanib or 100nM MEKi in combination with 500nM MCL1i and stratified by niche states: high, mixed, and low cancer proximities.

**Figure 7.**
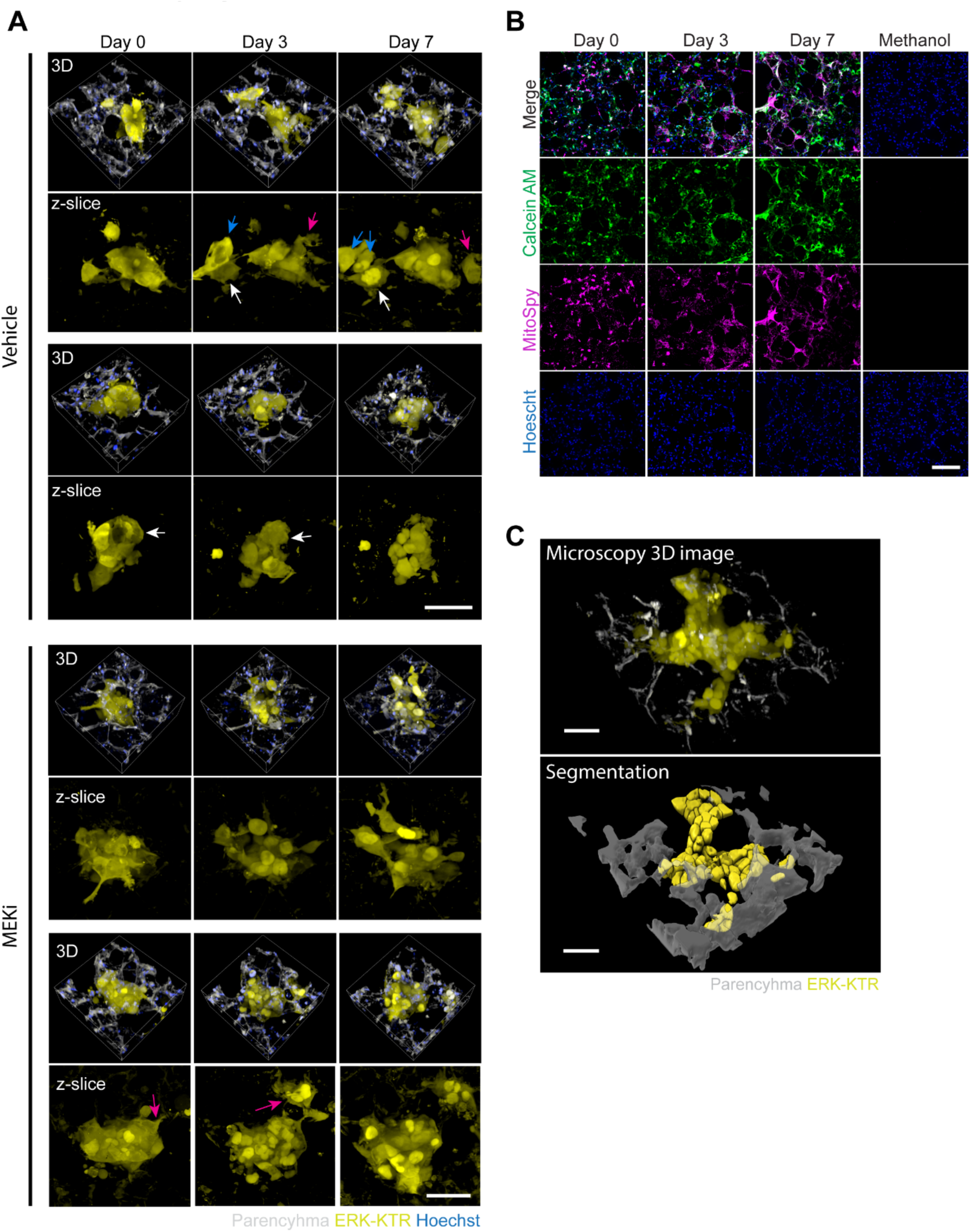
Comparison of ex vivo and in vivo drug response. **(A)** Images (left) showing vimentin-stained OS-17 metastases in lung sections obtained from mice bearing three-week metastasis and treated for 7 days in vivo with vehicle, MCL1i, pazopanib, or a combination of MCL1i and pazopanib (see methods for dosages). Images show vimentin stain (green) and lung parenchyma (grey). Corresponding masks obtained from NIS-Elements software are shown on the right. Scale bars = 200 µm. **(B)** Images (left) showing OS-17 reporter cells in LungSITE after 7 days of treatment with vehicle, 100nM MCL1i, 2µM pazopanib or a combination of MCL1i and pazopanib ex vivo. Images show ERK-KTR (yellow) and lung parenchyma (grey). Corresponding masks obtained from SITE computational pipeline are shown on the right. Scale bars = 20 µm. **(C)** Violin plots comparing colony areas of individual OS-17 metastases measured from vimentin staining of lung sections obtained from in vivo treatment of mice, or reporter presence in lung slices treated ex vivo (LungSITE). Means and standard error of mean are indicated by a black circle and error bars, respectively. Lesions with colony areas bellow 300 µm were excluded. LungSITE data was obtained from 3 replicate experiments, while in vivo data was obtained from a study with 40 mice, with ≥5 mice per condition **(D)** Significance heatmap showing p-values from t-tests between in vivo and LungSITE colony areas used in (C). Both methods produced comparable overall metastasis size distributions across conditions, with only minimal significant differences between corresponding in vivo and LungSITE groups. In both models, MCL1i and pazopanib monotherapy reduced metastasis size relative to vehicle, whereas the combination treatment produced the greatest effect. Although the overall response patterns were highly concordant between the two models, statistical significance of individual comparisons varied, likely reflecting differences in sample size between the in vivo and LungSITE experiments. These findings show that LungSITE model captures key features of in vivo drug response and would be a valuable preclinical tool for evaluating drug efficacy.

## SUPPLEMENTARY TABLES

**Supplementary Table 1.**
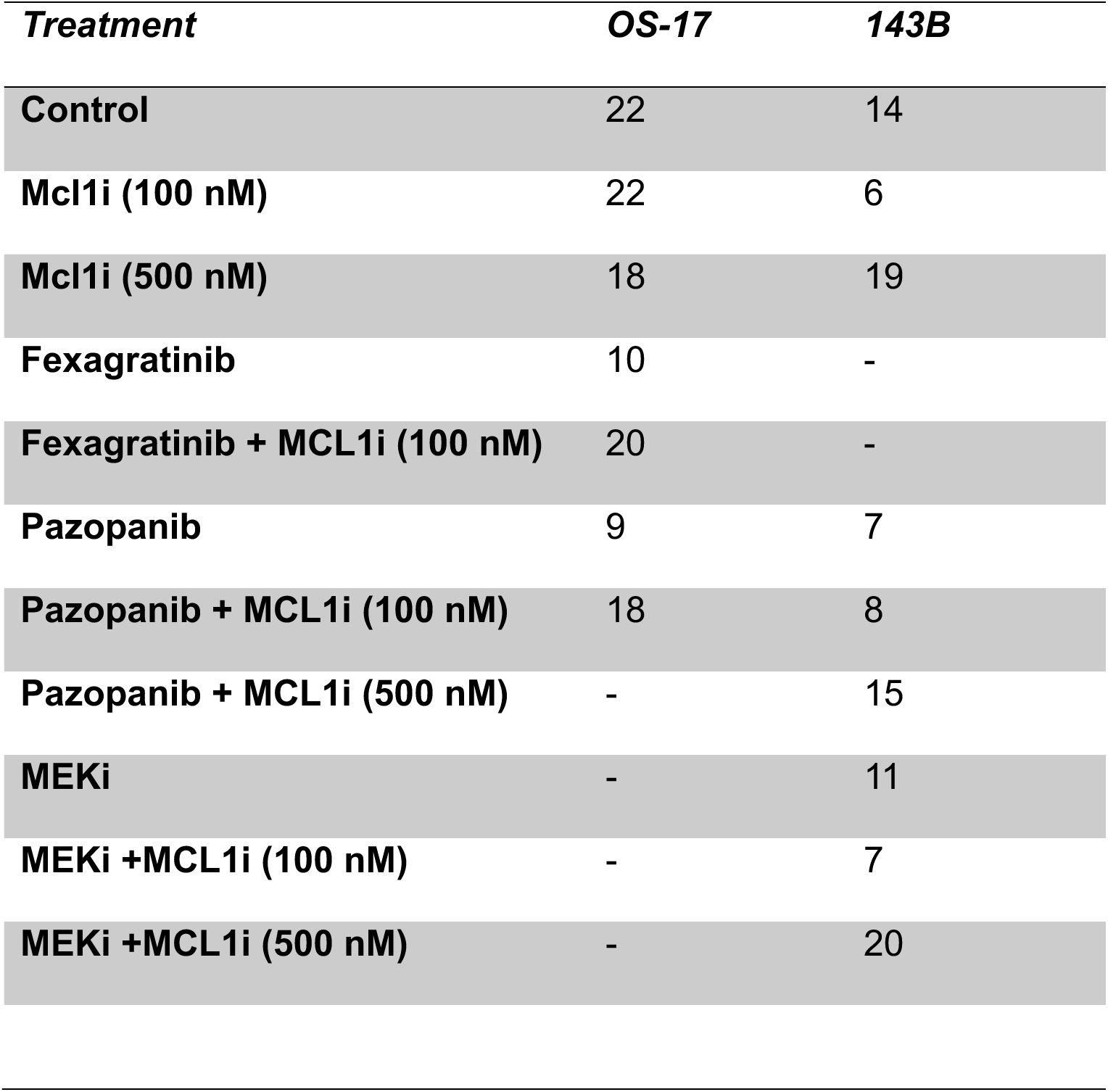
Summary of imaging datasets used for analysis, outlining number of XYs analyzed per treatment condition for each cell line.

**Supplementary Table 2.** Summary of total number of mice injected to obtain PCLS per cell line.

| **Cell line** | **OS-17** | **143B** |
| --- | --- | --- |
| **Number of injections** | 7 | 2 |

